# The obstacles and potential clues of prime editing applications in tomato, a dicot plant

**DOI:** 10.1101/2021.03.15.435378

**Authors:** Tien Van Vu, Jihae Kim, Swati Das, Jinsu Lee, Jae-Yean Kim

## Abstract

Precision genome editing is highly desired for crop improvement. The recently emerged CRISPR/Cas technology offers great potential applications in precision plant genome engineering. A prime editing (PE) approach combining a reverse transcriptase (RT) with a Cas9 nickase and a “priming” extended guide RNA has shown a high frequency for precise genome modification in mammalian cells and several plant species. Nevertheless, the applications of the PE approach in dicot plants are still limited and inefficient. We designed and tested prime editors for precision editing of a synthetic sequence in a transient assay and for desirable alleles of 10 loci in tomato by stable transformation. Our data obtained by targeted deep sequencing also revealed only low PE efficiencies in both the tobacco and tomato systems. Further assessment of the activities of the PE components uncovered potential reasons for the inefficiency of the PE complexes. Modifying the pegRNA sequences by shortening or introducing mismatches to the primer binding sequences (PBS) in order to reduce their melting temperatures (Tm) did not enhance the PE efficiency at the SlBMP21, SlALC and SlALS1 loci. Our data show challenges of PE approach in tomato, indicating a further improvement of the PE system for the successful applications such as use of improved expression systems. Our work provides an important clue for the successful application of the PE approach in crop improvement.

## INTRODUCTION

RNAs have been used for gene correction in human and yeast cells (Meers et al., 2016). CRISPR/Cas-mediated precision editing using a chimeric single guide RNA (sgRNA) and a donor template was shown in rice protoplasts (Butt et al., 2017; Li et al., 2019). However, the gene targeting (GT) frequency recorded using the RNA donor was much lower than that of a single-stranded DNA (ssDNA) donor (Li et al., 2019). Further work needs to be done on the use of RNA as a template for precision plant gene editing.

Recently, prime editing (PE) using guide RNA extensions for priming reverse transcription-mediated precise editing appeared to be an excellent precision genome editing technique in mammalian cell lines (Anzalone et al., 2019). The best version of the prime editor used a CRISPR/Cas complex developed by fusing a reverse transcriptase (RT) to the C-terminus of a Cas9 nickase (H840A) and a PE gRNA (pegRNA) with a 3’ extension that could bind to the 3’ nicked strands produced by nCas9. When bound, the nicked strand’s free 3’-OH is used as the substrate for the RT to copy genetic information from the pegRNA’s 3’ extension. If pegRNAs were designed to produce modified nucleotides, these nucleotides would be inserted into the genome during downstream repair processes (Anzalone et al., 2019). A second nick site present downstream of the first nick site would support the retention of the introduced nucleotides. The PE approach may also be an excellent alternative to GT with shorter editing sequence coverage (Van Vu et al., 2019). Prime editors have been shown to work well in monocot plants by Lin and coworkers (Lin et al., 2020). In the same year, other reports also showed the activities of PE complexes in monocots such as rice (Butt et al., 2020; Hua et al., 2020; Li et al., 2020; Tang et al., 2020; Xu et al., 2020) and maize (Jiang et al., 2020). However, the majority of the data showed successful PE at the acetolactate synthase (ALS) locus. Very limited data regarding the applications of PE in dicots were released, with relatively low efficacy in potato (Veillet et al., 2020) and tomato (Lu et al., 2020), and this process would need to be improved for further applications.

Therefore, we sought to investigate the activity of PE complexes in plants using a transient assay with a synthetic substrate in tobacco and stable transformation for editing ten loci in tomato. Our data showed low PE efficiency at both the somatic cell and plant levels. Additional analyses revealed the possible negative effects of the PE components on PE performance. Our work shows both obstacles and important clues for the further improvement of the PE approach in plants.

## RESULTS AND DISCUSSION

### The PE complexes did not perform better than the BE tool in a transient assay

The PE tool includes an nCas9 fused with an RT enzyme and a pegRNA for binding to the targeted sequence, and its 3’ extension primes an RNA-dependent DNA polymerization reaction by annealing to the sequence located upstream of the nicked site on the nontargeted strand (Figure 1a). If a second nick is simultaneously added, we have a PE3 or PE3b approach, depending on the position of the second nick outside or inside the area covered by the RT, respectively (Anzalone et al., 2019). To quickly validate the PE activity in plants, we started our PE experiments with a transient *Agrobacterium*-mediated infiltration system in tobacco leaves that was designed to modify a single nucleotide of a synthetic substrate ((CA)n substrate) (Figure 1b and 1c). A cytidine base editor (CBE) system using PmCDA1 cytosine deaminase was used for comparison of the editing efficiency. The CBE design included a “-sgRNA” control (pCEC1) and two test vectors, one with the T-DNA (pCE01) and one with a geminiviral replicon system (pCE02) (Vu et al., 2020), for amplifying the editing tools (Figure 1c). The PE3b tools were designed with both T-DNA and replicon cargos but also included dual vector systems to avoid possible activity in bacterial cells. One vector carried the nCas9-RT expression cassette, the synthetic substrate, and a sgRNA expression cassette for generating a second nick (pPEsubc1 and pPEsubc2), and the other vector contained the pegRNA expression cassette (pPEsub1 and pPEsub2) (Figure 1b and 1c). RT DNA was chemically synthesized as a tobacco codon-optimized Moloney murine leukemia virus (M-MLV) RT based on the PE2 approach used in the work of Anzalone and coworkers (Anzalone et al., 2019). pPEsubc1 and pPEsubc2 were also used alone as controls for the PE experiments. Three temperature conditions were applied to the infiltrated plants during the incubation stage to study the impacts of temperature on the activities of the BE and PE complexes. Surprisingly, we observed a very strong C to T transition at the 19^th^ nucleotide counted from the PAM by BE in *Agrobacterium* cells (Figure 1d and 1e, *Agrobacteria* panel) that led to indistinguishable BE activity at that position in tobacco leaves (Figure 1d and 1e). By contrast, we observed BE activities in converting C to T at the 15^th^ and 17^th^ nucleotides in the tobacco leaves (Figure 1d and 1e). Interestingly, the BE activities were reduced when the temperature increased from 25°C to 37°C (Figure 1e and 1f and Supplemental Table 2). Our data also indicate that the replicon-based BE tool worked much more efficiently than the T-DNA tool under transient expression conditions (Figure 1e and 1f and Supplemental Table 2). Unexpectedly, we failed to show any PE activity with the transient system using both the T-DNA and replicon systems. Furthermore, no improvement was obtained using the temperature treatments (Figure 1g and 1h). One of the reasons for the failure of the PE3b constructs might be due to the inefficiency of the dual *Agrobacterium*-mediated infiltration that required simultaneous transfer of the two T-DNAs carrying PE components into one cell.

**Figure 1.**
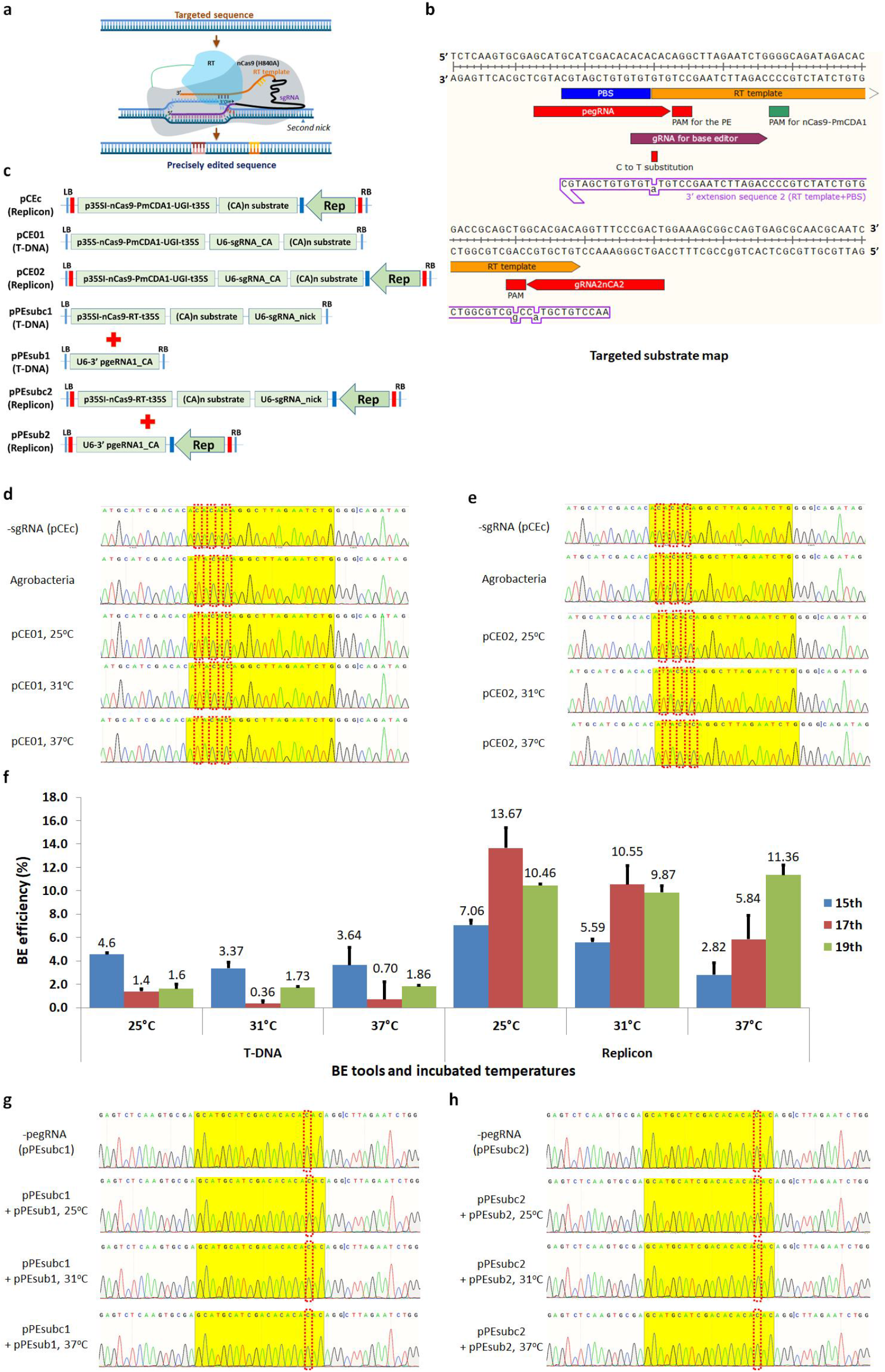
Failure of prime editing in tobacco cells in transient assays. **(a)** Schematic processes of primed editing. The nontargeted strand is nicked by CRISPR/Cas9 nickase (H840A), and the 3’ nicked end binds to a complementary RNA (priming binding site, PBS, 13-15 nt) introduced by extending the 3’ end of the sgRNA (RT template) and primes reverse transcription by the nCas9-fused RT using its free OH group. RT copies the genetic information from the RNA template that is complementary to the sequence downstream from the nicked site and includes intentionally introduced base modifications (indicated by red and bright orange lines). The RT product appears as a 3’-flap sequence that competes with the full complement of the original 5’-flap during the repair process, and a precisely edited sequence could be fixed to the targeted site after 5’-flap removal and subsequent replication of the DNA. A second nick is introduced during the improvement of the prime editor for supporting 3’-edited flap fixation. **(b)** Selected synthetic target and pegRNA design. pegRNA and its 3’ extension and second nick spacer sequences are indicated. A PmCDA1-based C->T editing tool is designed in parallel for comparison, and its spacer sequence is also shown. Intended base modifications are denoted and explained. **(c)** Vector arrangements for the study. The nCas9(D10A)-PmCDA1 base editing system is designed with the control plasmid (pCEc, -gRNA), T-DNA (pCE1), and replicon (pCE2)-based editors. The prime editors are designed with a dual vector system for T-DNA (pPEsubc1 + pPEsub1) and replicon (pPEsubc2 + pPEsub2)-based tools for editing C->T and C->A (shown in **b**). **(d-e)** Sequencing data showing C->T editing by T-DNA- (**d**) and replicon-based (**e**) tools (Supplemental Table 2, (**e**)). The –sgRNA (pCEc) chromatogram represents all the temperature tested. **(f)** Editing efficiency of the base editors. The T-DNA construct represents the pCE1 plasmid, and the replicon construct is for the pCE2 plasmid. The editing efficiencies at different temperatures are plotted for the 15^th^, 17^th^, and 19^th^ bases counted from the PAM site to its upstream spacer sequence. **(g-h)** Failure of the T-DNA- (**g**) and replicon-based (**g**) prime editors to edit the intended bases. The -pegRNA (pPEsubc1) and -pegRNA (pPEsubc2) chromatograms represent for all the temperature tested. Discontinuous red boxes denote the intended modifications in each of the editing conditions (25, 31, and 37°C) for the editors and the control.

### The PE tools showed low precise editing efficiencies at 10 tomato loci

We decided to evaluate the ability of the PE system to edit genomic sites in tomato by Agrobacterium-mediated stable transformations. The tomato transformation system was efficiently used for GT experiments in our lab (Supplemental Figure 1) (Vu et al., 2020). We first attempted to edit three tomato loci, SlHKT1;2, SlEPSPS1, and SlOr using PE2 and PE3 approaches with the T-DNA and replicon systems (Supplemental Figure 2 and 3 and Data S1). Transformed explants collected at 10 days post-transformation (dpt) were subjected to targeted deep sequencing of the targeted sites. However, in two replicates, we could not reveal any evidence of PE activity that was higher than the background level of the targeted sequencing method (Table 1). PE activity was shown to be locus-specific, and several parameters, such as PBS and RT template lengths, may also affect PE efficiency in plants (Lin et al., 2020). During our study, several PE data were reported that showed comparable PE efficiencies among the PE2, PE3, and PE3b constructs (Butt et al., 2020; Hua et al., 2020; Lin et al., 2020; Tang et al., 2020), and in some cases, the PE2 approach performed better than the PE3 approach (Jiang et al., 2020; Veillet et al., 2020). We then extended the PE experiment to 7 more loci using the PE2 approach and single replicon-based vectors (Figure 2a and Supplemental Table 3) to check whether the PE system would work in tomato. Most of the loci were selected to require only single nucleotide changes, and the RT template lengths were designed within the optimal ranges shown in reports on monocots. The PBS lengths were also appropriately selected (Table 1). Using targeted deep sequencing analysis, we found a higher PE performance at SlWH9, SlKD1, SlALC, and SlALS1 than at the mock control, albeit at low absolute values (Table 1). The low PE efficiency might be partially explained by the sample type (cotyledon explants) we collected for targeted deep sequencing. Since the cotyledon explants (0.1 × 0.3 cm in size) contained many untransformed cells due to the nature of the Agrobacterium-mediated transformation, they confer much lower editing efficiency than the protoplast system. Therefore, we also analyzed plants regenerated from the transformed cotyledon explants. We detected only low levels of PE alleles in several plants transformed with the SlALC and SlALS1 PE constructs (Figure 2c and d). Sanger sequencing data revealed that only 1% of the PE alleles were successfully fixed in the plant genomes. Nevertheless, the PE efficiency obtained at the plant stage in our work is comparable to that of recently published data in tomato (Lu et al., 2020). Moreover, in one event at SlALC, two byproducts were also formed, with frequencies of 1 and 3% (Figure 2d).

**Table 1.** Prime editing efficiency in tomato using tobacco codon-optimized RT and rice codon-optimized PPE.

| No. | Locus | Targeted change | PBS length (nt) | RT length (nt) | Second nick position | Mock control |  | nCas9-RT |  | nCas9-PPE |  |
| --- | --- | --- | --- | --- | --- | --- | --- | --- | --- | --- | --- |
|  |  |  |  |  |  | Total reads | PE-mediated precise editing efficiency (%) | Total reads | PE-mediated precise editing efficiency (%) | Total reads | PE-mediated precise editing efficiency (%) |
| 1 | SIHKT1;2 | N217D: +21 T to C; +10 A to G (reverse complement) | 12 | 26 | +76 (PE3) | 321183 | 0.007 | 104610 | 0.000 | - | - |
|  |  |  | 12 | 26 | PE2 |  |  | 151412 | 0.000 | 312118 | 0.000 |
| 2 | SIEPSPS1 | T178I: +16 TG to GA; P182S: +4 CGG to AGA (reverse complement) | 15 | 20 | -55 (PE3) | 2526 | 0.000 | 1357 | 0.000 | - | - |
|  |  |  | 15 | 20 | PE2 |  |  | 1380 | 0.000 | 351540 | 0.000 |
| 3 | SIOr | R95H: +1 AGG to CAT | 12 | 14 | +61 (PE3) | 449607 | 0.000 | 140325 | 0.000 | - | - |
|  |  |  | 12 | 14 | PE2 |  |  | 155826 | 0.000 | 322070 | 0.000 |
| 4 | SIMBP21 | Premature Q108stop: +1 C to T | 13 | 13 | PE2 | 49811 | 0.018 | 57943 | 0.022 | 148426 | 0.019 |
| 5 | WH9 | G69D (s-classic): +6 G to A | 13 | 14 | PE2 | 59484 | 0.018 | 54735 | 0.110 | 202599 | 0.032 |
| 6 | SIKD1 | 1 bp deletion at 1266 bp upstream of an open reading frame (ORF): +21 A del | 13 | 26 | PE2 | 44911 | 0.002 | 60765 | 0.018 | 169327 | 0.008 |
| 7 | SIPRD | A46 deletion: DELLAVLG to DELLVLG: +18 CTG del | 13 | 22 | PE2 | 46727 | 0.000 | 50399 | 0.000 | 130662 | 0.000 |
| 8 | SIALC | ALC to alc allele: T to A; V106D: +2 A to T (reverse complement) | 13 | 12 | PE2 | 51754 | 0.015 | 45543 | 0.042 | 159889 | 0.064 |
| 9 | SIDMR6-1 | Premature truncation of the protein; E140stop: +1 G to T | 13 | 13 | PE2 | 37532 | 0.128 | 50231 | 0.020 | 170289 | 0.021 |
| 10 | SIALS1 | P186S: +1 C to T | 13 | 13 | PE2 | 56496 | 0.057 | 63128 | 0.095 | 201599 | 0.095 |

**Figure 2.**
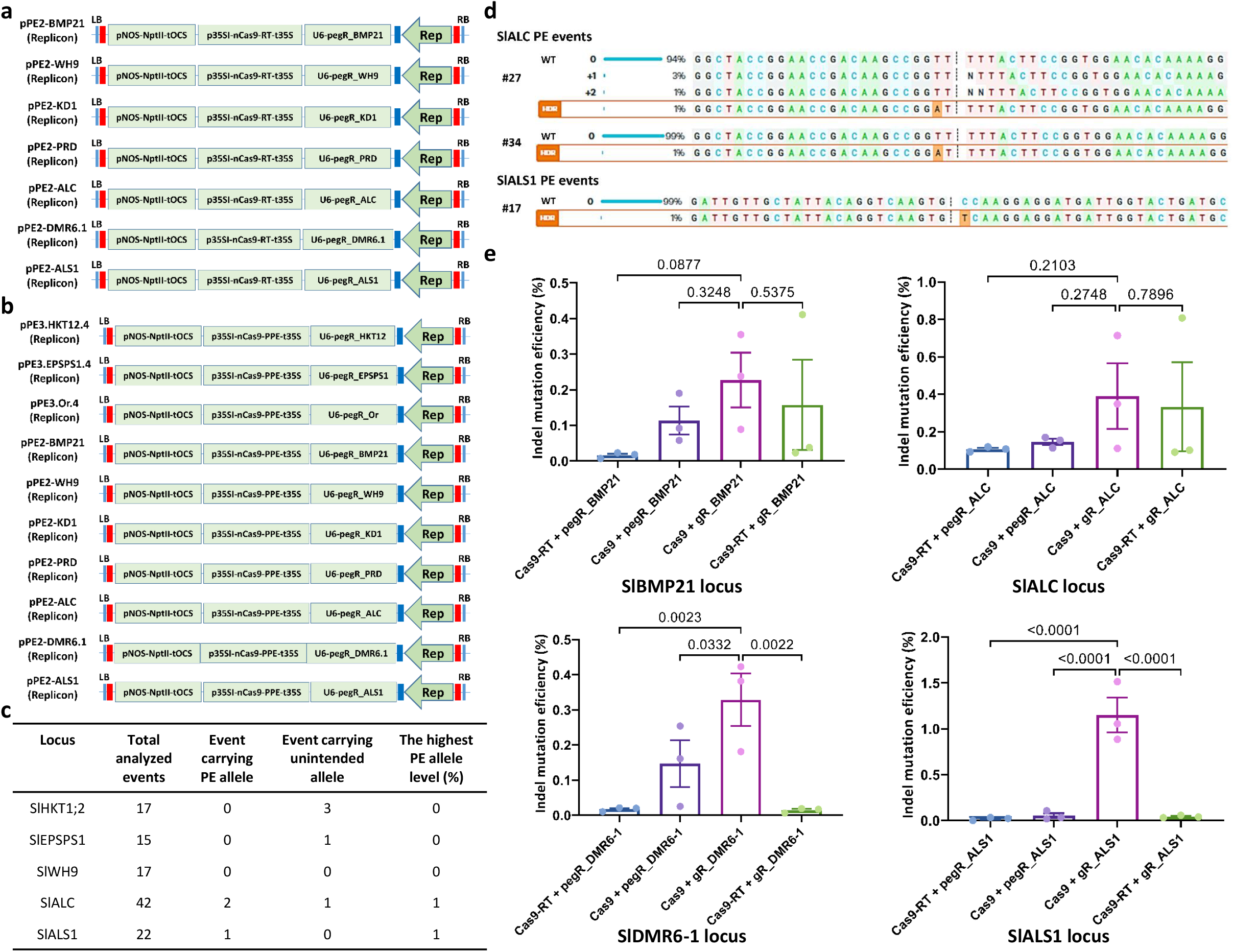
Prime editing performance in tomato. **(a-b)**. PE2 prime editing constructs using the geminiviral replicon in tomato with synthetic tobacco codon-optimized RT (**a**, 7 loci) or rice codon-optimized PPE (**b**, 10 loci). The NptII expression cassette (Addgene plasmid #51144) was used for the kanamycin selection of transformed cells, tissues, and plants. The expression of the prime editors was driven by the p35S promoter with the UBQ10 intron 1 and CaMV 35S terminator (Data S1). The transcription of the pegRNAs was controlled by the AtU6 core promoter sequence (Data S1). **(c)** PE performance data obtained from the analysis of the plant-stage transformants of SlWH9, SlALC, and SlALS1 by Sanger sequencing and ICE Synthego. **(d)** The traces of PE-edited alleles were obtained for SlALC (upper two panels) and SlALS1 (bottom panel). The precisely edited alleles are shown at the bottom of each panel and are denoted by “HDR” labels. The PE-mediated replaced bases are highlighted with an orange background. **(e)** Scatter dot-bar plots showing the pairwise comparisons between the indel mutation efficiencies of the Cas9-gRNA complexes with the PE components and that of the normal construction at the SlBMP21, SlALC, SlDMR6-1, and SlALS1. The comparison at each locus is presented with the locus name at the bottom of the plot. The combinations (denoted as “+”) of nuclease complexes are shown at the bottom of each scattered bar with Cas proteins (Cas9 and Cas9-RT fusion) and guide RNAs (normal gRNA with gR prefix and pegRNA with pegR prefix, followed by the locus name). The sequences flanking the targeted sites were amplified at 10 dpt and analyzed by targeted deep sequencing in three replicates. The indel mutation efficiencies were revealed by RGEN Cas-Analyzer. Multiple comparisons of the means and plotting were conducted by GraphPad Prism version 9 using uncorrected Fisher’s LSD test. The p-values of each compared mean pair are shown on the top of the bars.

The very low PE efficiency obtained for 10 tomato loci based on various editing types under our experimental conditions indicates that many critical factors affected the overall performance of PE in tomato and other dicot plants, as also concluded previously (Veillet et al., 2020). To exclude the possibility that our tobacco codon-optimized RT coding sequence may trigger aberrant transcription or cryptic splicing, thereby affecting nCas9-RT activity, next, we used the nCas9-PPE coding sequence that showed good PE activities in rice and wheat (Lin et al., 2020). The nCas9-PPE prime editor is a rice codon-optimized RT that performed the best among the RTs tested by Lin and coworkers. We constructed PE2 tools using nCas9-PPE with all 10 tomato loci tested earlier (Figure 2b). However, targeted deep sequencing data collected for two replicates showed similar levels of PE activity across all 10 loci. The PE efficiency was even reduced in the case of the SlWH9 and SlKD1 loci but improved at the SlALC locus (Table 1), indicating that our tobacco codon-optimized RT activity was not worse than the reported version. These data illustrate that the PE approach needs improvement for further applications in tomato.

### The PE components negatively affect dsDNA cleavage

In an attempt to find solutions for improving the PE performance, we sought to investigate the roles of the PE components (i.e., the nCas9-RT and the pegRNAs) in the activation of the PE nuclease complex and nicking the template strand. We constructed binary plasmids with fully functional SpCas9 fused with RT or SpCas9 alone in combination with either pegRNAs or normal sgRNAs for four loci (SlBMP21, SlALC, SlDMR6-1, and SlALS1) (Supplemental Figure 4) and assessed their ability to generate indel mutations by targeted deep sequencing. Unexpectedly, the Cas9-RT fusion combined with the pegRNAs did not show indel mutation efficiencies above background levels at three (SlBPM21, SlDMR6-1, and SlALS1) out of the four loci (Figure 2e), while the conventional Cas9-sgRNA complexes worked well at all the tested loci and significantly higher at the SlDMR6-1 and SlALS1 loci (Figure 2e and Supplemental Table 4). Cas9-RT combined with normal sgRNAs generated indel mutations at 2 of the 4 tested loci (SlBMP21 and SlALC), with indel mutation efficiencies similar to those of Cas9 and normal gRNAs (Figure 2e and Supplemental Table 4). The pegRNAs also supported the formation of indel mutations when combined with the Cas9 nuclease at 3 out of the 4 tested loci (SlBMP21, SlALC, and SlDMR6-1), but the indel mutation efficiencies were much low compared to that of the Cas9/sgRNA complexes (Figure 2e and Supplemental Table 4). Either of the PE components alone worked at extremely lower efficiencies compared to Cas9 with normal gRNA at the ALS1 locus. These data indicate that the PE components negatively affected the cleaving activity of the Cas9 nuclease and that the use of pegRNAs led to reduced activity of the Cas9 enzyme (Figure 2e). We reason that the Cas9-RT fusion might reduce the accessibility of the larger active complexes (Cas9-RT is approximately 236 kDa, compared to ∼158 kDa for SpCas9 alone) to some genomic contexts, such as SlDMR6-1 and SlALS1, thereby blocking their cleavage activity. Moreover, the pegRNAs shared intramolecular complementarity between PBS and spacer sequences that potentially altered the secondary structure of the sgRNAs (Supplemental Figure 5), leading to possible interference of SpCas9 activation and/or the ability of gRNA to bind to targeted sequences and cleave them. The failure of the Cas9-RT/pegRNA complexes to produce indel mutations at three out four loci and very low efficiency at the other locus might have resulted from the combined negative impacts of both components. Based on these data, we hypothesize that the activity of nCas9-RT/pegRNA complexes was affected by the negative impacts of both RT fusion and pegRNA structure, which led to extremely low PE efficiency, and by some unknown factor, resulting in especially low PE efficiencies in tomato and other dicot plants (Lu et al., 2020; Veillet et al., 2020).

### Modifications of the PBS sequences did not significantly improve PE efficiency at three tested loci

If the above assumption is true, then we expect that when the PBS is shortened, the PE efficiency will be improved. However, the PE data in plants suggested optimal ranges of PBS lengths that could not be too short to prime RT polymerization (Lin et al., 2020). Higher incubation temperatures may help to destabilize the PBS-spacer intramolecular interactions, but they may also affect the activity of nuclease complexes or the activity of its target binding, as observed with the transient BE data (Figure 1). That is also why Lu and coworkers could not obtain a significant improvement in PE activity when they incubated explants at high temperatures (Lu et al., 2020). During the preparation of this manuscript, Lin and coworkers reported that the optimal melting temperature (Tm) of the PBS sequence was around 30°C for PE in rice (Lin et al., 2021) reasoning that the PBS Tm affects the priming activity. Thus, we hypothesize that a range of optimized melting temperature exists for an active PBS of pegRNA in tomato. Alternatively, introducing several mismatches between the PBS and spacer sequences may significantly weaken the interaction and improve PE activity in plants.

To test the hypothesis, we modified the pegRNAs of the three loci (SlBMP21, SlALC, and SlALS1) by introducing mismatches to the original or shortened PBS sequences (Fig. 3a). Total four different modifications to an original PBS were made and binary plasmids with replicon were similarly constructed (Fig. 3a, b) to the original PE vectors (Fig. 2a) using the tobacco codon-optimized RT. The PE efficiency was assessed by targeted deep sequencing. In three replicates, the PE efficiencies at all the three tested loci were not significantly improved compared to that of the original pegRNAs (Fig. 3c and Supplemental Table 5). These data were revealed even with the modified PBS with Tm 30, 32, or 34°C compared to the original 38°C Tm (Fig. 3c and Supplemental Table 5). It is still unknown if the way of introducing the mismatches was not perfectly matched with the criteria of the PBS Tm optimization in the case of the data shown by Lin and coworkers (Lin et al., 2021) or the mismatch itself was not sufficient to improve the PE performance in tomato.

**Figure 3.**
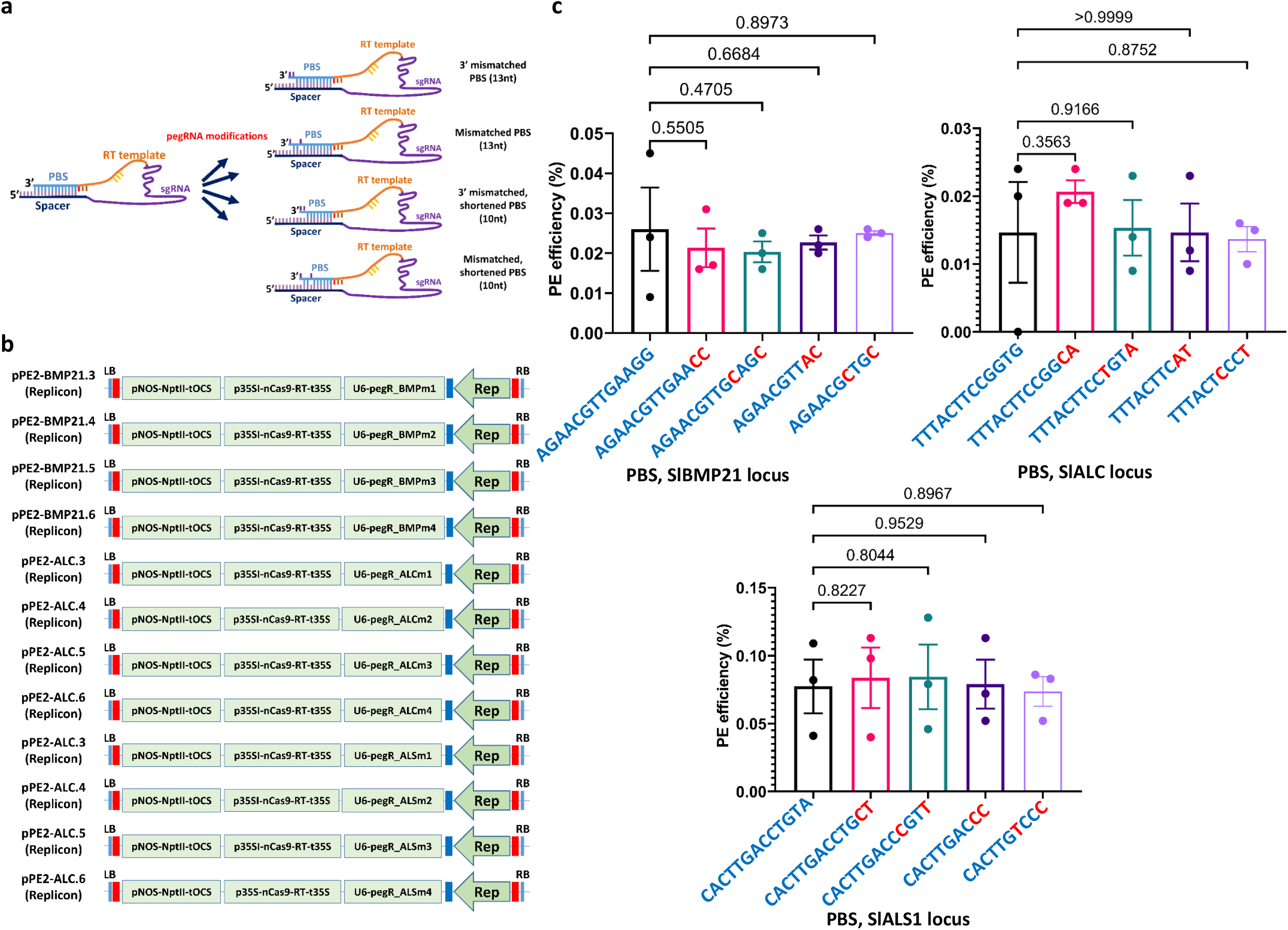
Assessment of prime editing efficiency using modified PBS sequences. **(a)** Modifications of the previously tested pegRNAs by shortened and/or introducing mismatches to their PBS sequences. There are four modifications that introduce mismatches to the original PBS sequences (13 nt in length, pegR.BMPm1, pegR.BMPm2, pegR.ALCm1, pegR.ALCm2, pegR.ALS1m1 and pegR.ALS1m2 for the BMP21, ALC and ALS1) or shortened PBS (10nt, pegR.BMPm3, pegR.BMPm4, pegR.ALCm3, pegR.ALCm4, pegR.ALS1m3 and pegR.ALS1m4 for the BMP21, ALC and ALS1) with mismatches. **(b)** PE2 prime editing constructs using the modified pegRNAs. **(c)** Scatter dot-bar plots showing the pairwise comparisons between the PE efficiencies with the original and modified pegRNAs at the SlBMP21, SlALC, and SlALS1. The comparison at each locus is presented with the locus name at the bottom of the plot. The PBS sequences are shown on the bottom of each bar plot with the red font letters represent mismatches between the PBS and spacer sequences. The sequences flanking the targeted sites were amplified at 10 dpt and analyzed by targeted deep sequencing in three replicates. The PE efficiencies were revealed by RGEN Cas-Analyzer. Multiple comparisons of the means and plotting were conducted by GraphPad Prism version 9 using uncorrected Fisher’s LSD test. The p-values of each compared mean pair are shown on the top of the bars.

Since both the pegRNA and Cas9-RT fusions affected the cleaving activity of the complex, we can also use a two-component PE complex that separates the nCas9 and RT polypeptides to reduce the space constraint caused by the nCas9-RT fusion. However, the two-component system may pose another risk for the proper assembly of the two components in a targeted genomic context. Taken together, the PE approach offers an excellent alternative to precision gene editing in plants. However, we could not obtain a better editing efficiency than that of the BE tools in a transient assay in tobacco. Similar results were also recorded for 10 tomato loci using stable *Agrobacterium*-mediated transformations. We revealed low but comparable PE efficiency at the plant stage compared with recent data in plants. Further analyses showed that the PE components (nCas9-RT fusion and pegRNA) might largely contribute to its low activity in tomato. However, our PBS sequence modifications for reducing Tm were failed to improve the PE efficiency. Significant improvement to the PE approach needs to be made for further applications in tomato, and our data shed light on potential solutions for the improvement of this editing tool.

## METHODS

### System design for PE experiments in tobacco and tomato

For the transient experiment, PE constructs were designed to modify two nucleotides of the (CA)n substrate sequence using the 3’ extension pegRNA (Figure 1a and 1b; Supplemental Data S1). A tobacco codon-optimized RT DNA sequence was fused with a human codon-optimized SpCas9 nickase (H840A) (nCas9-RT, Supplemental data S1) cloned using SpCas9 of the Addgene plasmid (#49771) that worked well in plants (Belhaj et al., 2013). The PE experiment was performed by using dual agroinfiltration (Figure 1c). For comparison, CBE constructs were designed to convert Cs to Ts, including the PE-targeted C at position 17 counted from the PAM of the CBE gRNA (Figure 1b).

For PE of the selected tomato loci using *Agrobacterium*-mediated stable transformation (Supplemental Figure 1), PE2 and PE3b approaches were applied for SlHKT1;2, SlEPSPS1, and SlOr (Supplemental Figures 2 and 3). Further applications of the nCas9-RT (tobacco codon-optimized RT (PE2) (Anzalone et al., 2019)), nCas9-PPE (rice codon-optimized RT (Lin et al., 2020)) systems using the original or modified PBS sequences on the other seven loci (Supplemental Table 3) used a PE2 strategy (Figure 2a and 2b; Supplemental Data S1).

In all binary vectors, the transcription of both CBE and nCas9-RT was driven by a long CaMV35S inserted with a UBQ10 intron 1 (Vu et al., 2020). The CBE gRNA and PE pegRNA were cloned downstream of an AtU6 promoter (Belhaj et al., 2013). The CBE and PE systems were delivered by either T-DNAs or geminiviral replicons (Vu et al., 2020).

### *Agrobacterium*-mediated infiltration and sample analysis

Leaves of tobacco (*Nicotiana benthamiana*) at the 6-leaf stage were infiltrated with *Agrobacteria* containing binary plasmids. *Agrobacteria* (GV3101::pMP90 strain) were prepared by an overnight primary culture with a subsequent 6-8 h secondary culture using liquid LB medium containing 75 mg/L kanamycin, 30 mg/L rifampicin, and 30 mg/L streptomycin. *Agrobacterium* cells were collected by centrifugation at 4,000 rpm for 10 minutes, resuspended in MS buffer containing 100 µM acetosyringone to an OD_600nm_ of 0.8, and incubated for one hour before infiltration. At three days post-infiltration, the infiltrated zones were cut, and total genomic DNA (gDNA) was isolated using the CTAB method. PCRs were then conducted using primers flanking the targeted sequences (Supplemental Table 1), and the gDNA templates and PCR products were purified and sequenced by the Sanger method. The editing efficiency was calculated based on the level of peaks of C and T at the targeted positions revealed by the chromatograms (Figure 1 d, 1e, 1g, and 1h).

### *Agrobacterium*-mediated tomato transformation and assessment of PE efficiency

*Agrobacterium*-mediated tomato transformation was performed using our in-house protocol (Supplemental Figure 1) (Vu et al., 2020). Briefly, seven-day-old cotyledons were cut and used for transformation. For targeted deep sequencing, thin cotyledon explants (0.1 × 0.3 cm) were used to reduce the area containing untransformed cells beyond the cut edges of the explants. The *Agrobacteria* containing the PE constructs were cultured and harvested as in the transient assay but were resuspended in an ABM-MS solution (Vu et al., 2020) containing 100 µM acetosyringone to an OD_600nm_ of 0.8. The *Agrobacteria* were then incubated for one hour before the transformation. Regenerated shoots were selected on a medium containing 80 mg/L kanamycin. Subsequently, hardened plants were checked for PE activity.

For the assessment of PE efficiency, samples were collected at 10 dpt and analyzed by targeted deep sequencing. At the plant stage, plant leaves were collected and screened for PE alleles by Sanger sequencing, and targeted deep sequencing analysis was conducted for the events carrying PE alleles.

### Targeted deep sequencing

Genomic DNA was isolated from cotyledon explants or plant leaves using the CTAB method. The MiniSeq sequencing service (MiniSeqTM System, Illumina, USA) was used. MiniSeq samples were prepared in three PCRs according to the manufacturer’s guidelines, with genomic DNAs as templates for the first PCR. The first and second PCRs used the primers listed in Supplemental Table 1, whereas the third PCR was conducted with the manufacturer’s primers to assign sample IDs. The MiniSeq raw data FASTQ files were analyzed by the Cas-Analyzer tool (Park et al., 2016) and PE-Analyzer (http://www.rgenome.net/pe-analyzer). The PE efficiency was assessed using the RT template sequences as the input HDR donor sequence.

### Data analysis and presentation

Most of the experiments analyzed by targeted deep sequencing were conducted in two to three replicates. The editing data, statistical analysis if applicable, and scattered plots were further processed by the MS Excel and Graphpad Prism 9.0 program. The multiple comparisons were conducted with the uncorrected Fisher’s LSD test. The data are appropriately explained in detail in the legends of figures and/or tables.

## ACKNOWLEDGMENTS

We wish to thank Mrs. Ngan Nguyen Thi for their valuable technical support in this study. This work was supported by the National Research Foundation of Korea (the Bio & Medical Technology Development Program 2020M3A9I4038352, 2020R1A6A1A03044344, 2020R1I1A1A01072130) and the Program for New Plant Breeding Techniques (NBT, grant PJ01478401), Rural Development Administration (RDA), Republic of Korea.

## AUTHOR CONTRIBUTIONS

T.V.V. and J.Y.K. conceived and designed the research. T.V.V., J.K., and S.D. conducted the experiments. T.V.V. and J.Y.K. analyzed the data. T.V.V. wrote the manuscript. T.V.V. and J.Y.K. finalized the manuscript. All authors read and approved the manuscript.

## SHORT LEGENDS FOR SUPPORTING INFORMATION

### List of Supplemental data file

Supplemental data S1

## Supplemental Figures, Tables and References

**Supplemental Figure 1:**
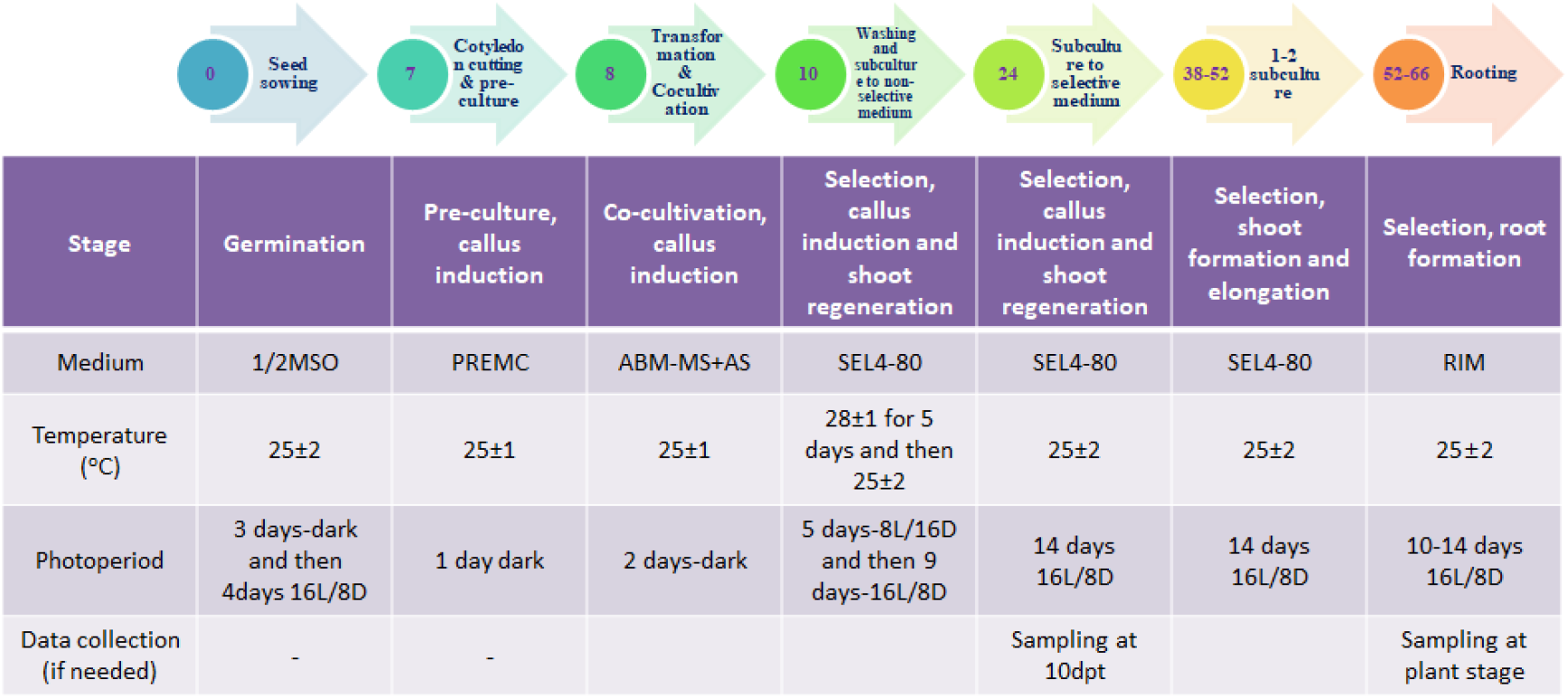
*Agrobacterium*-mediated transformation protocol used in this work. The step-by-step protocol is presented with each number in the circles indicating the number of days after seed sowing (upper panel), and the treatments used in each step are shown in the lower panel.

**Supplemental Figure 2:**
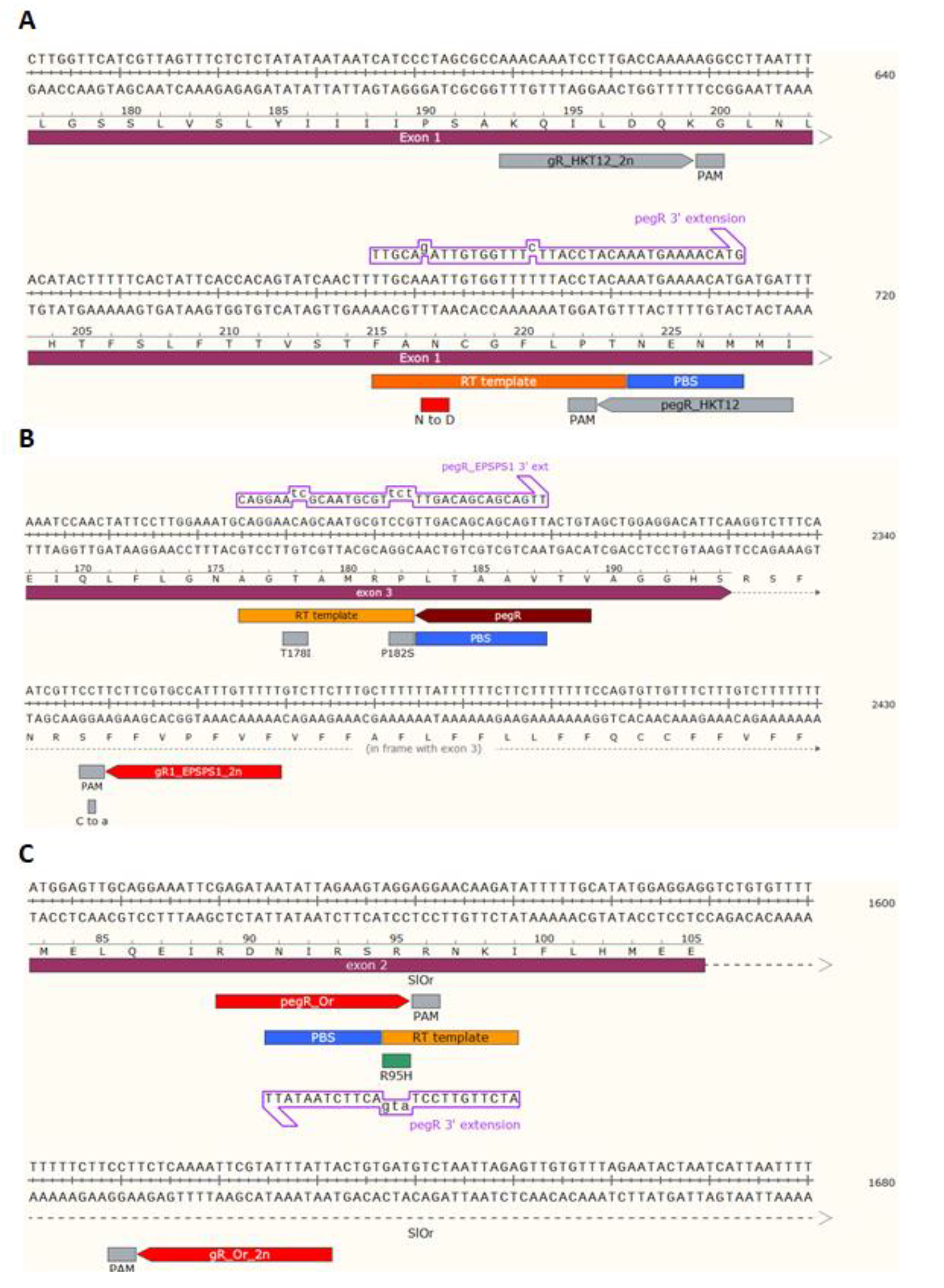
PE targeting maps of the first three loci. (A) SlHKT1;2. (B) SlEPSPS1. (C) SlOr.

**Supplemental Figure 3:**
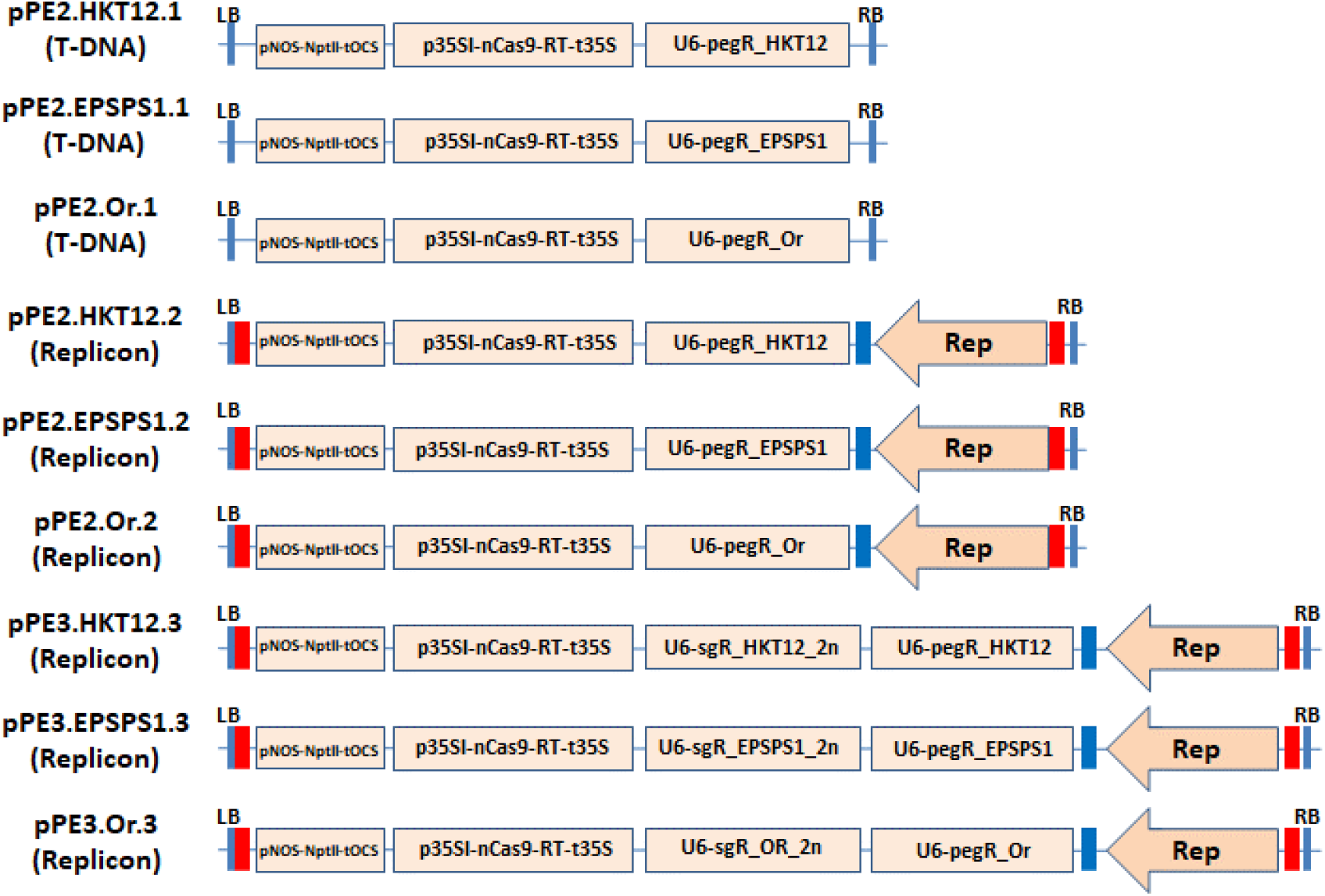
PE constructs for editing the SlHKT1;2, SlEPSPS1 and SlOr loci by PE2 and PE3 approaches.

**Supplemental Figure 4:**
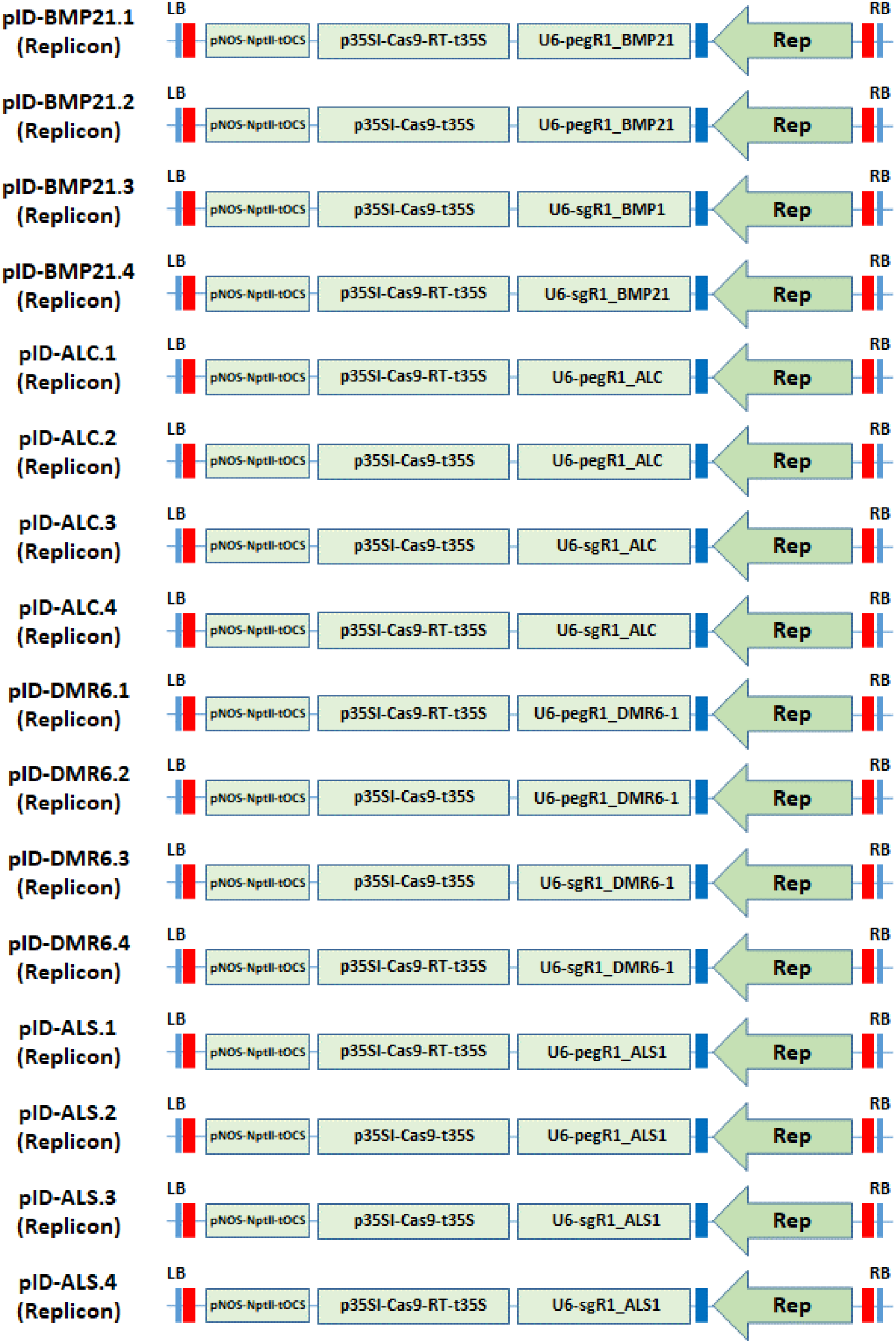
Constructs used for the assessment of indel mutation formation by different combinations of either Cas9-RT or Cas9 with either sgRNAs or pegRNAs at the SlBMP21, SlALC, SlDMR6 and SlALS1 loci. All the component were expressed with the geminiviral replicon system.

**Supplemental Figure 5:**
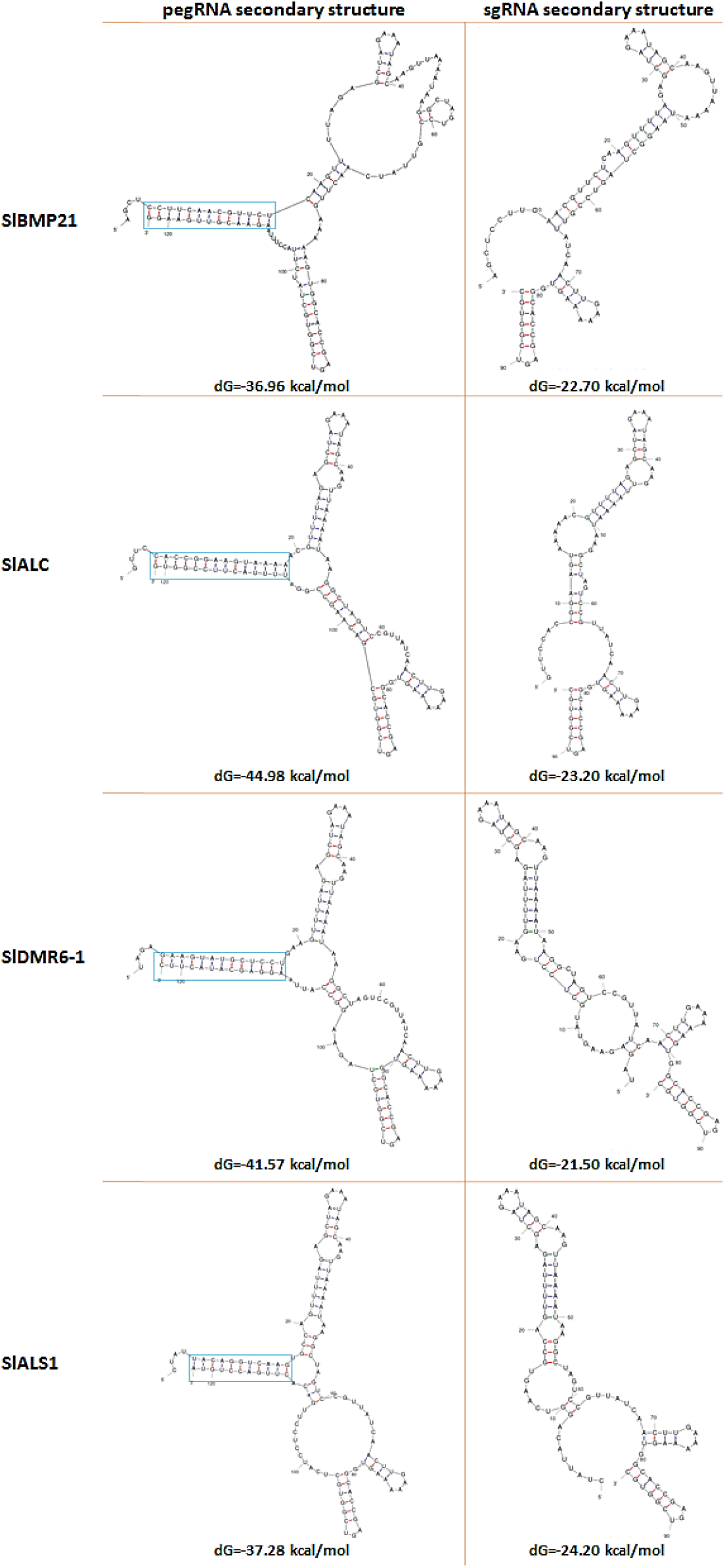
MFOLD-predicted secondary structures of pegRNAs and sgRNAs of SlBMP21, SlALC, SlDMR6-1 and SlALS1. The pegRNA structures (left panel) and sgRNA structures (right panel) were predicted using MFOLD software. The deltaG (dG) of each structure is denoted at the bottom of it. The discontinuous light blue boxes indicate the intermolecular annealing of the PBS and spacer sequences.

**Supplemental Table 1.**
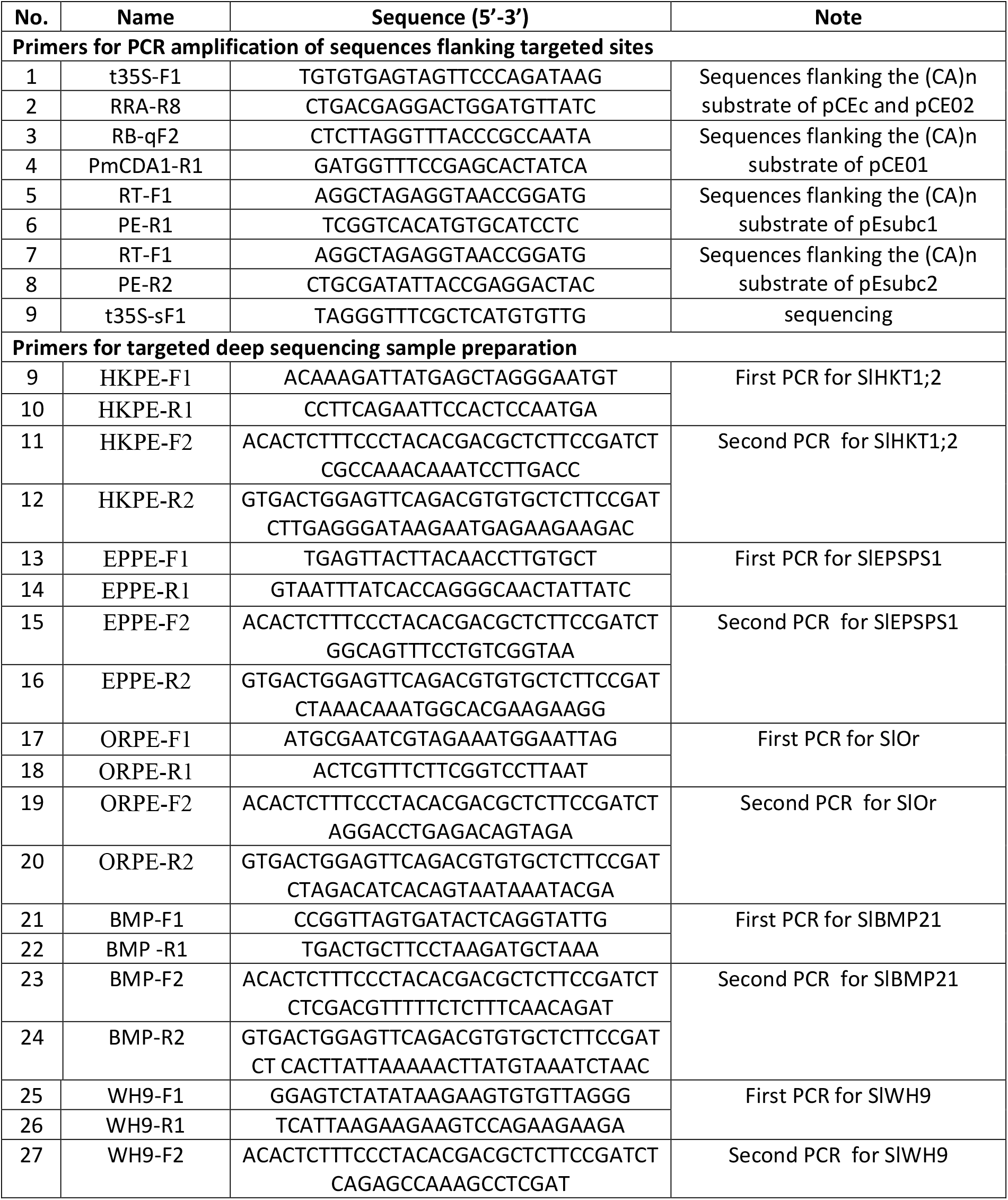
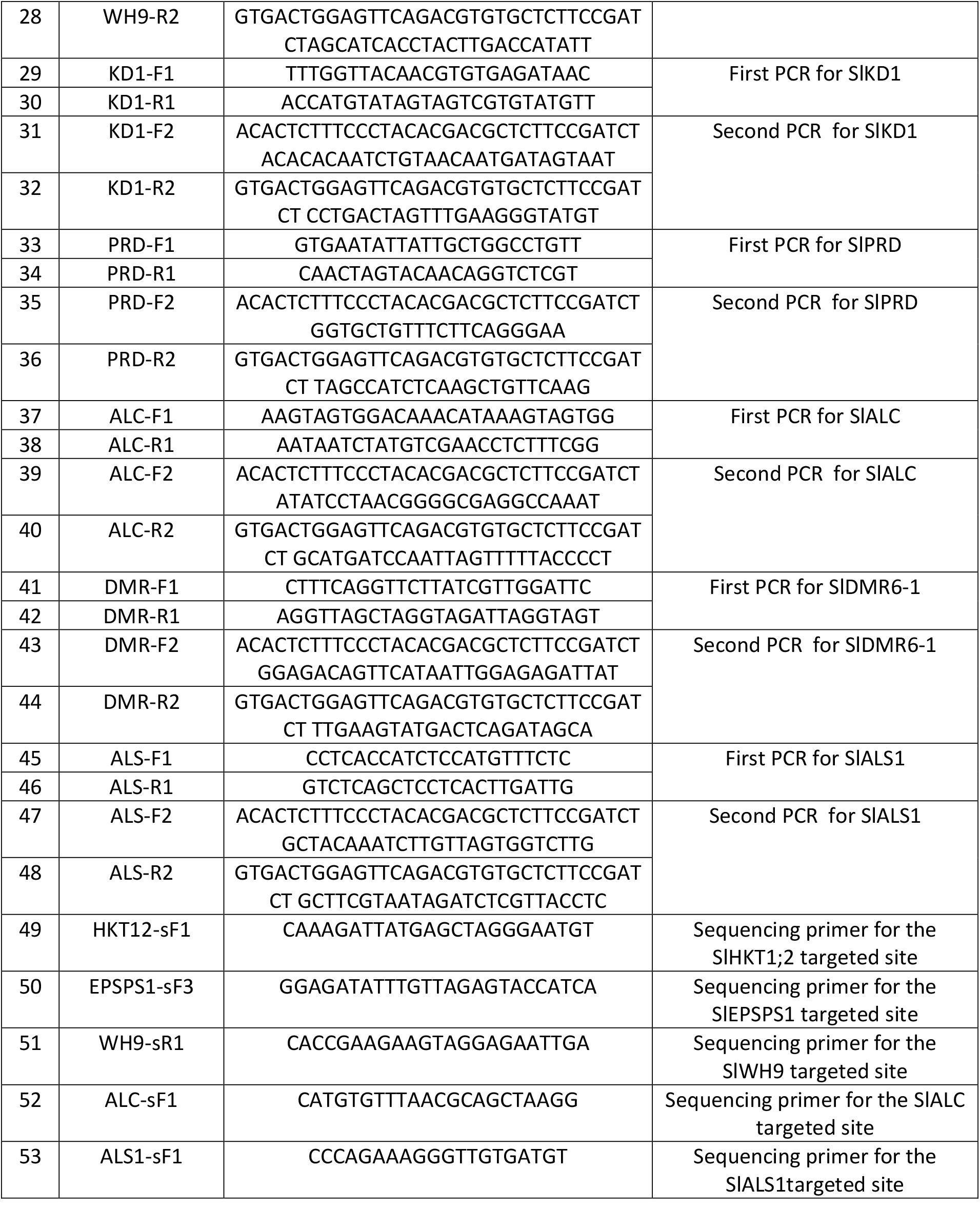
Sequences and primers used in this study.

**Supplemental Table 2.** Base editing efficiency obtained in the tobacco leaf infiltration assay.

| Position counted from PAM | T-DNA |  |  | Replicon |  |  |
| --- | --- | --- | --- | --- | --- | --- |
|  | 25°C | 31°C | 37°C | 25°C | 31°C | 37°C |
| 15th | 4.57 ± 0.14 | 3.37 ± 0.55 | 3.64 ± 1.5 | 7.06 ± 0.41 | 5.59 ± 0.25 | 2.82 ± 1.02 |
| 17th | 1.38 ± 0.21 | 0.36 ± 0.22 | 0.70 ± 1.5 | 13.67 ± 1.74 | 10.55 ± 1.63 | 5.84 ± 2.03 |
| 19th | 1.63 ± 0.41 | 1.73 ± 0.10 | 1.86 ± 0.03 | 10.46 ± 0.10 | 9.87 ± 0.57 | 11.36 ± 0.83 |

**Supplemental Table 3.**
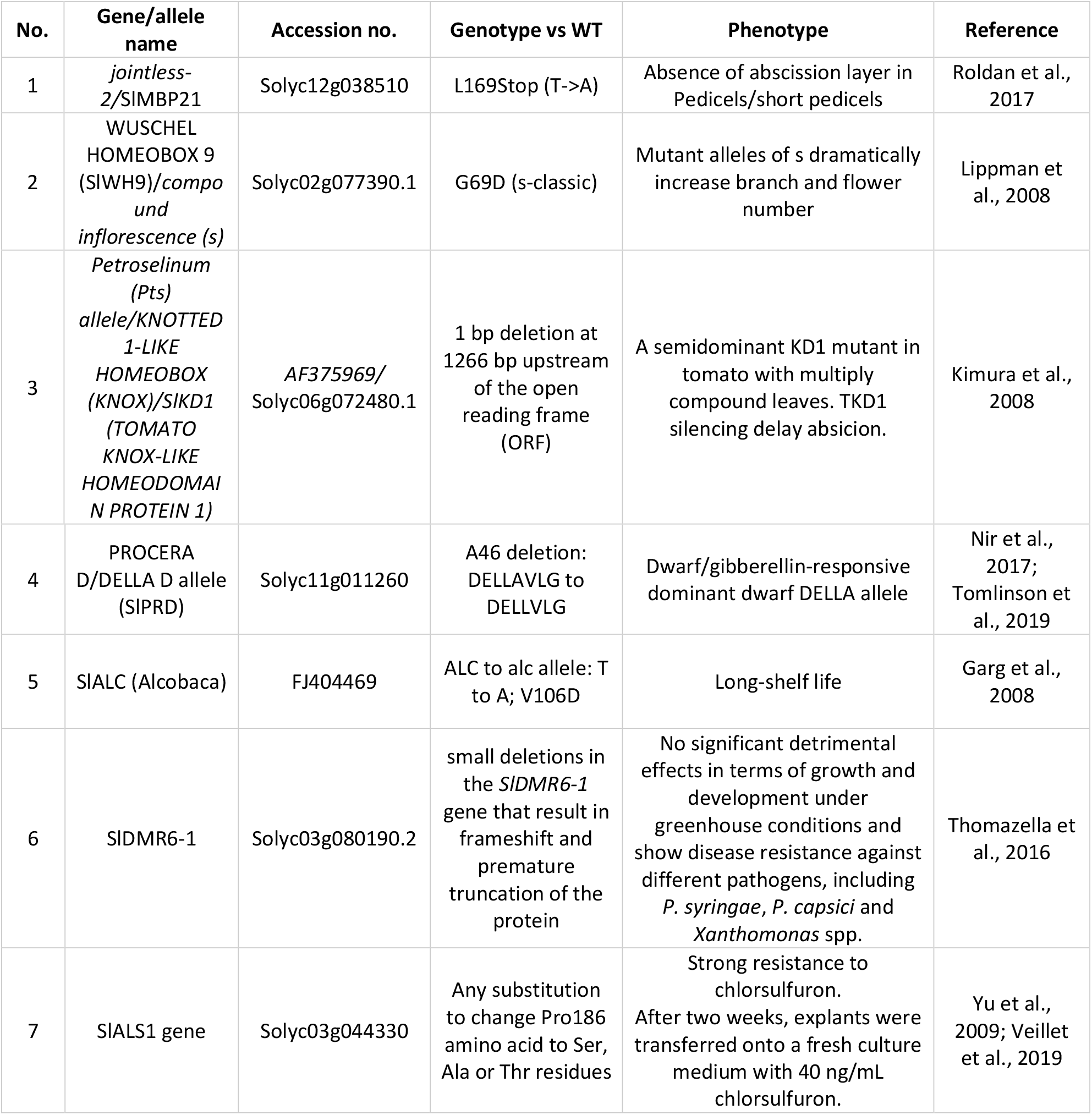
PE targets in tomato used in the study.

| No. | Gene/allele name | Accession no. | Genotype vs WT | Phenotype | Reference |
| --- | --- | --- | --- | --- | --- |
| 1 | *jointless-2*/SIMBP21 | Solyc12g038510 | L169Stop (T->A) | Absence of abscission layer in Pedicels/short pedicels | Roldan et al., 2017 |
| 2 | WUSCHEL HOMEODOMAIN 9 (SIWH9)/*compound inflorescence (s)* | Solyc02g077390.1 | G69D (s-classic) | Mutant alleles of s dramatically increase branch and flower number | Lippman et al., 2008 |
| 3 | *Petroselinum* (Pts) allele/*KNOTTED 1-LIKE HOMEODOMAIN (KNOX)/SIKD1 (TOMATO KNOX-LIKE HOMEODOMAIN PROTEIN 1)* | AF375969/<br>Solyc06g072480.1 | 1 bp deletion at 1266 bp upstream of the open reading frame (ORF) | A semidominant KD1 mutant in tomato with multiply compound leaves. TKD1 silencing delay abscission. | Kimura et al., 2008 |
| 4 | PROCERA D/DELLA D allele (SIPRD) | Solyc11g011260 | A46 deletion: DELLAVLG to DELLVLG | Dwarf/gibberellin-responsive dominant dwarf DELLA allele | Nir et al., 2017;<br>Tomlinson et al., 2019 |
| 5 | SIALC (Alcobaca) | FJ404469 | ALC to alc allele: T to A; V106D | Long-shelf life | Garg et al., 2008 |
| 6 | SIDMR6-1 | Solyc03g080190.2 | small deletions in the *SIDMR6-1* gene that result in frameshift and premature truncation of the protein | No significant detrimental effects in terms of growth and development under greenhouse conditions and show disease resistance against different pathogens, including *P. syringae*, *P. capsici* and *Xanthomonas* spp. | Thomazella et al., 2016 |
| 7 | SIALS1 gene | Solyc03g044330 | Any substitution to change Pro186 amino acid to Ser, Ala or Thr residues | Strong resistance to chlorsulfuron. After two weeks, explants were transferred onto a fresh culture medium with 40 ng/mL chlorsulfuron. | Yu et al., 2009; Veillet et al., 2019 |

**Supplemental Table 4.** Comparisons of indel mutation efficiency of the Cas9 complexes containing the PE components compared to the normal complex.

| Gene/allele name | Accession no. | Construction | PBS length (nt) | RT length (nt) | Total reads | Indel mutation efficiency (%) |  |  |  |  | p value** (Uncorrected Fisher's LSD) |
| --- | --- | --- | --- | --- | --- | --- | --- | --- | --- | --- | --- |
|  |  |  |  |  |  | Replicate 1 | Replicate 2 | Replicate 3 | Average | SEM |  |
| SIBMP21 | Solyc12g038510 | -* | - | - | 49819 | 0.026 | - | - | 0.026 | - | - |
|  |  | Cas9-RT + pegR_BMP21 | 13 | 13 | 132955 | 0.018 | 0.007 | 0.023 | 0.016 | 0.005 | 0.0877 |
|  |  | Cas9 + pegR_BMP21 | 13 | 13 | 139784 | 0.092 | 0.190 | 0.058 | 0.113 | 0.040 | 0.3248 |
|  |  | Cas9 + gR_BMP21 | 0 | 0 | 127696 | 0.355 | 0.238 | 0.089 | 0.227 | 0.077 |  |
|  |  | Cas9-RT + gR_BMP21 | 0 | 0 | 128919 | 0.411 | 0.023 | 0.038 | 0.158 | 0.127 | 0.5375 |
| SIALC | FJ404469 | - | - | - | 51907 | 0.075 | - | - | 0.075 | - | - |
|  |  | Cas9-RT + pegR_ALC | 13 | 12 | 140946 | 0.119 | 0.108 | 0.093 | 0.107 | 0.007 | 0.2103 |
|  |  | Cas9 + pegR_ALC | 13 | 12 | 140852 | 0.170 | 0.155 | 0.114 | 0.146 | 0.017 | 0.2748 |
|  |  | Cas9 + gR_ALC | 0 | 0 | 129357 | 0.714 | 0.111 | 0.349 | 0.391 | 0.175 |  |
|  |  | Cas9-RT + gR_ALC | 0 | 0 | 121087 | 0.091 | 0.808 | 0.102 | 0.334 | 0.237 | 0.7896 |
| SIDMR6-1 | Solyc03g080190 | - | - | - | 37540 | 0.019 | - | - | 0.019 | - | - |
|  |  | Cas9-RT + pegR_DMR6-1 | 13 | 13 | 108382 | 0.010 | 0.021 | 0.020 | 0.017 | 0.004 | 0.0023 |
|  |  | Cas9 + pegR_DMR6-1 | 13 | 13 | 106969 | 0.254 | 0.161 | 0.025 | 0.147 | 0.067 | 0.0332 |
|  |  | Cas9 + gR_DMR6-1 | 0 | 0 | 124113 | 0.423 | 0.382 | 0.181 | 0.329 | 0.075 |  |
|  |  | Cas9-RT + gR_DMR6-1 | 0 | 0 | 106991 | 0.006 | 0.017 | 0.019 | 0.014 | 0.004 | 0.0022 |
| SIALS1 | Solyc03g080190 | - | - | - | 56500 | 0.000 | - | - | 0.000 | - | - |
|  |  | Cas9-RT + pegR_ALS1 | 13 | 13 | 131194 | 0.000 | 0.032 | 0.023 | 0.018 | 0.009 | <0.0001 |
|  |  | Cas9 + pegR_ALS1 | 13 | 13 | 145331 | 0.033 | 0.106 | 0.018 | 0.052 | 0.027 | <0.0001 |
|  |  | Cas9 + gR_ALS1 | 0 | 0 | 144979 | 1.055 | 1.513 | 0.884 | 1.151 | 0.188 |  |
|  |  | Cas9-RT + gR_ALS1 | 0 | 0 | 129222 | 0.039 | 0.057 | 0.031 | 0.042 | 0.008 | <0.0001 |

\*WT sample. \*\*Multiple comparisons were conducted between the normal Cas9+gRNA combination and the complex containing either or both the PE components with the Uncorrected Fisher's LSD test using the Graphpad 9.0 software.

**Supplemental Table 5.**
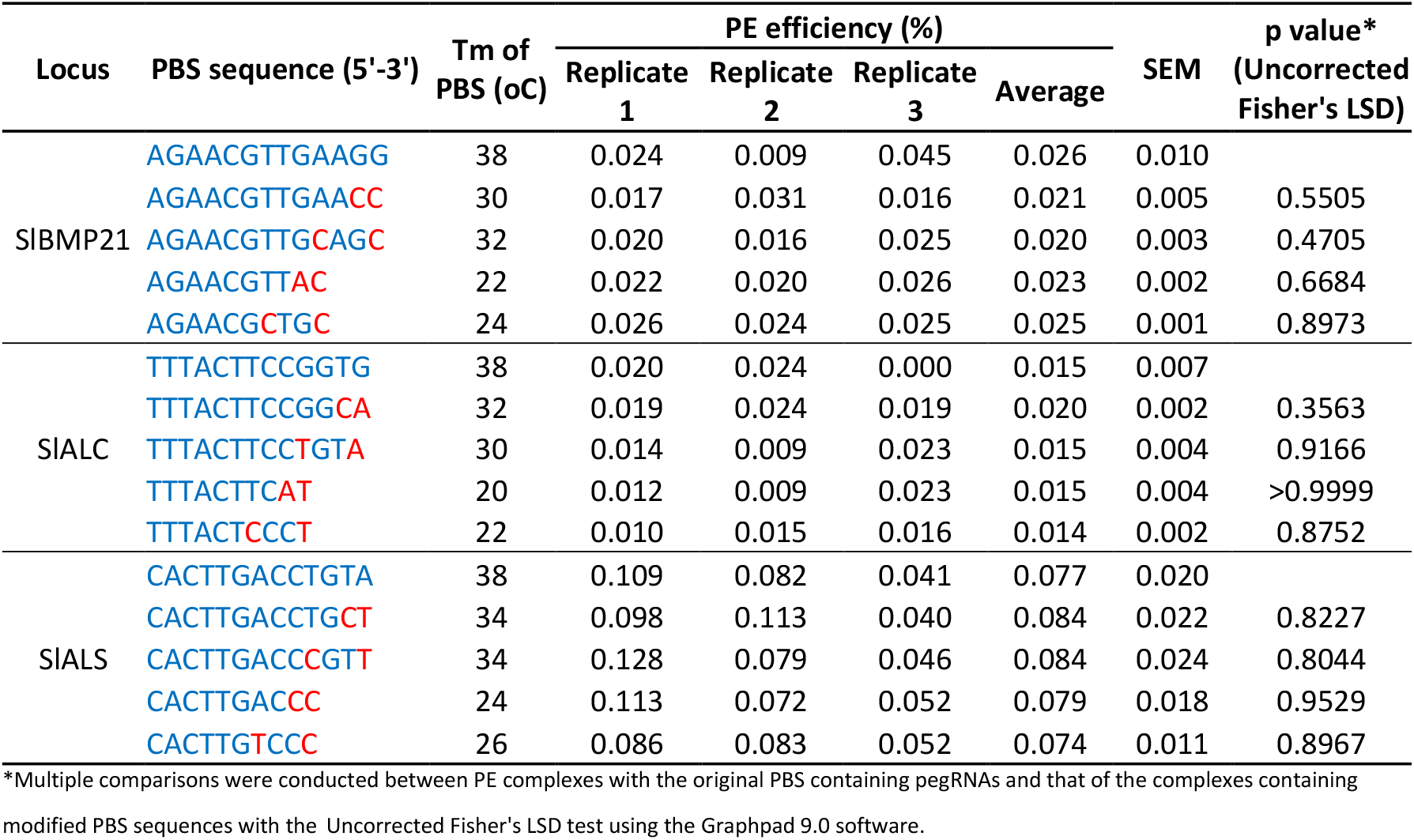
Comparisons of indel mutation efficiency of the Cas9 complexes containing the PE components compared to the normal complex.

## SEQUENCES USED IN THE STUDY

✥ **Moloney Murine Leukemia Virus reverse transcriptase (MMLVrt) coding sequence (tobacco codon-optimized)**

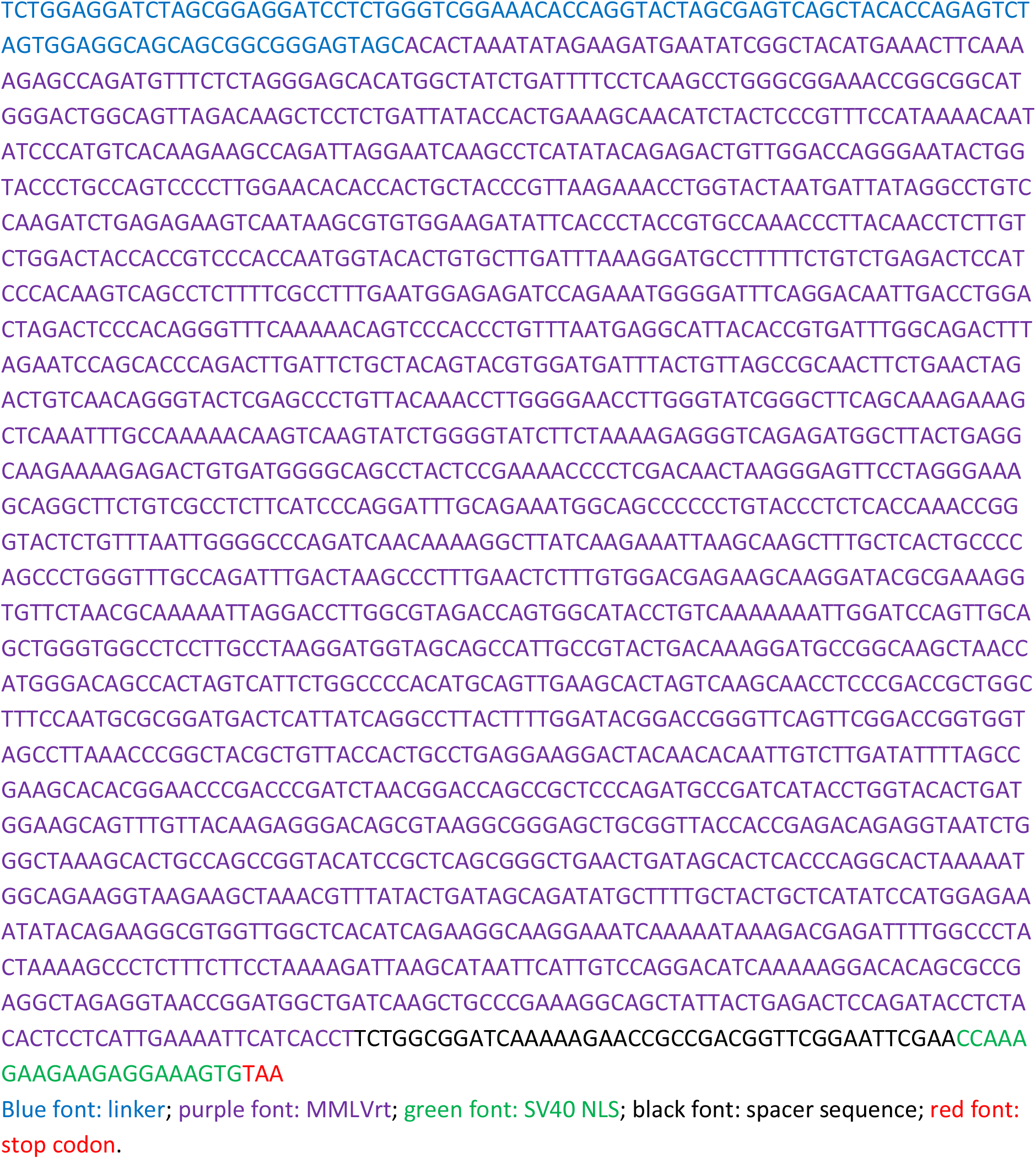
✥ **Moloney Murine Leukemia Virus reverse transcriptase (MMLVrt) coding sequence (rice codon-optimized) from Addgene Plasmid #140445**.
✥ **(CA)n substrate sequence**

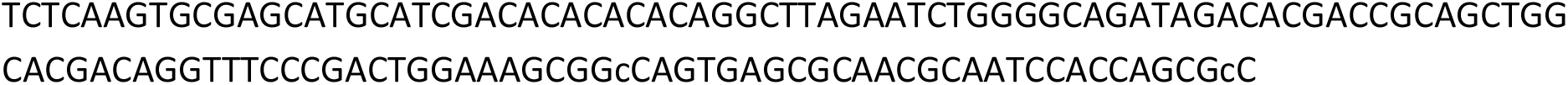
✥ **p35SI-nCas9-PmCDA1-UGI-t35S**

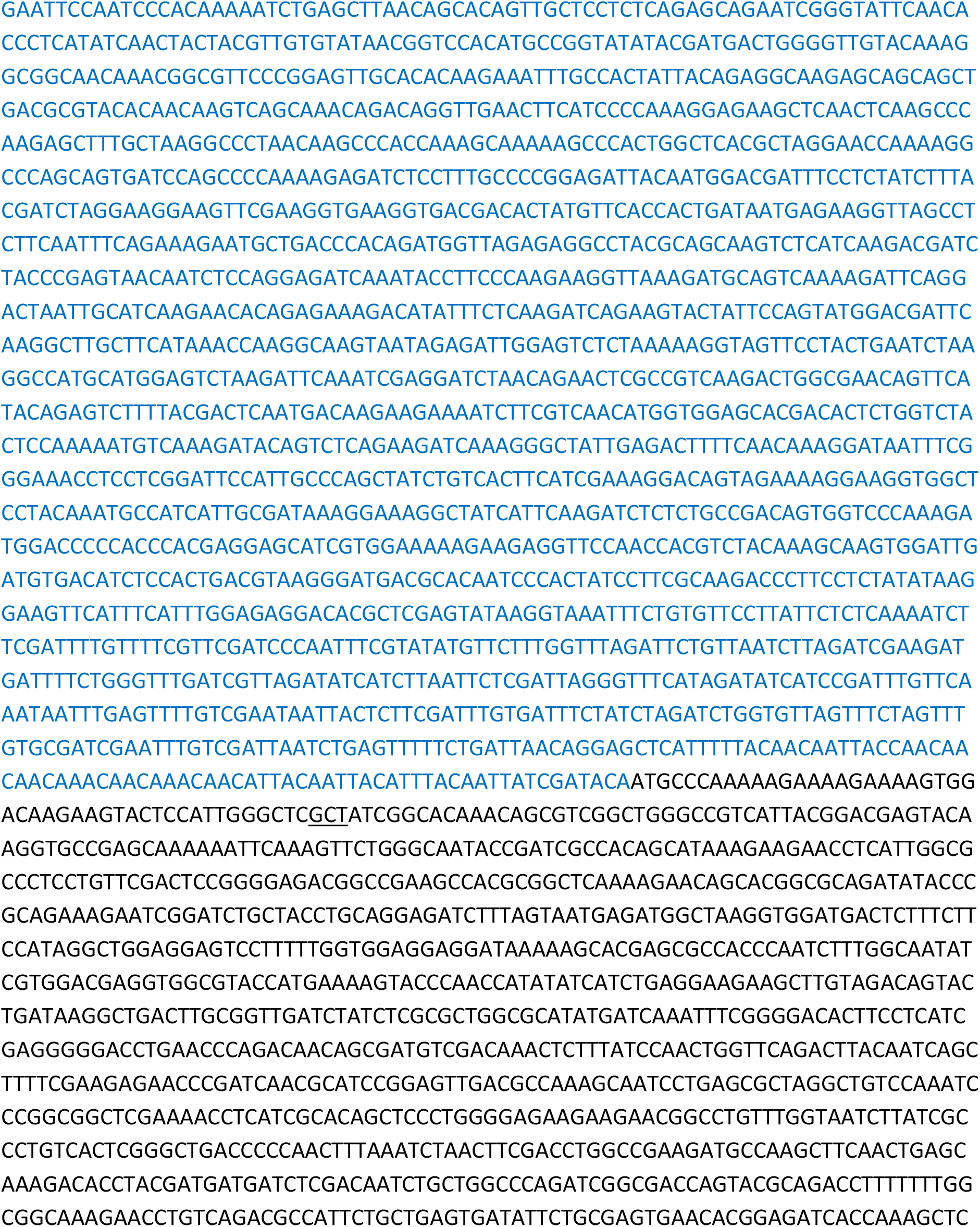

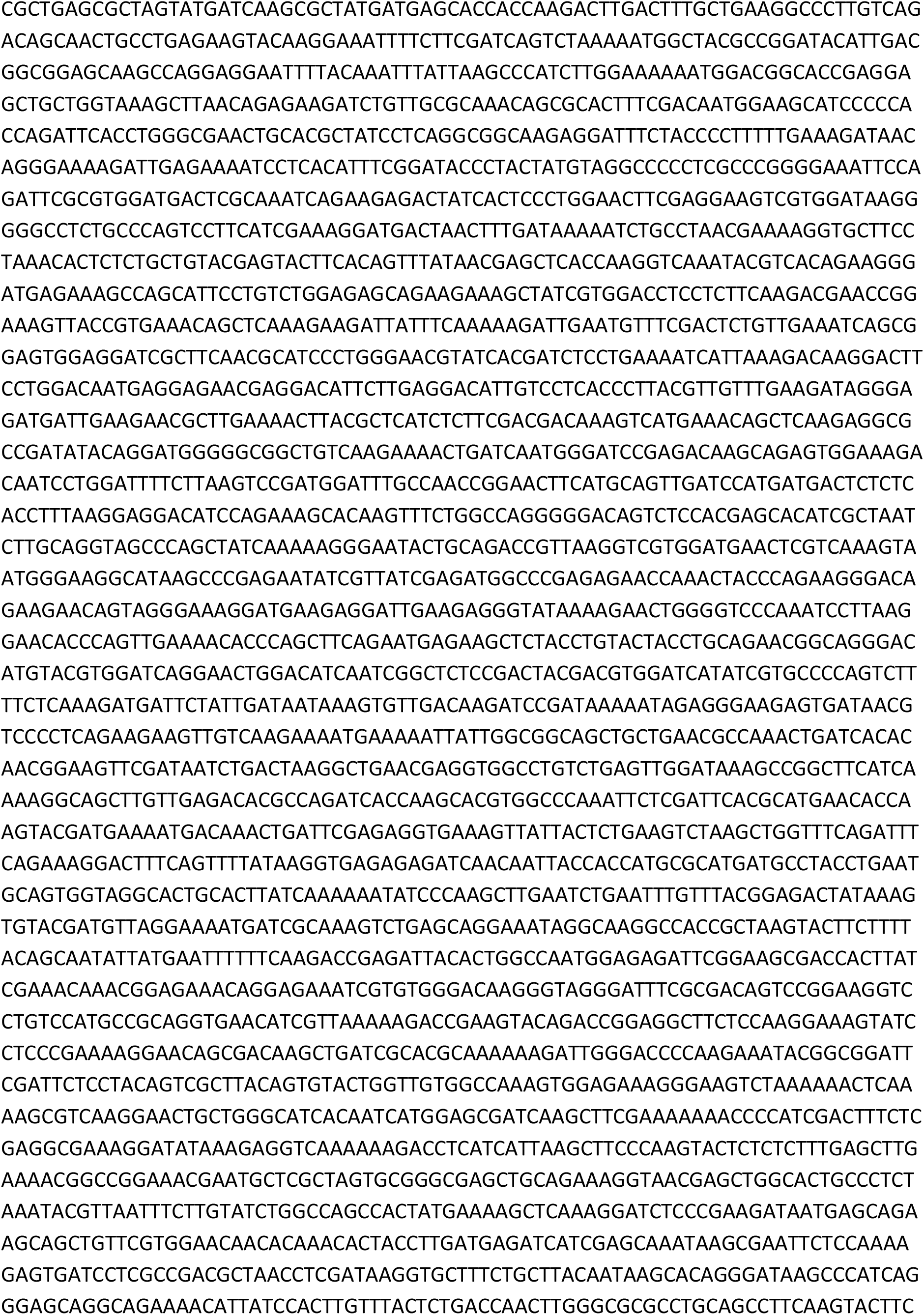

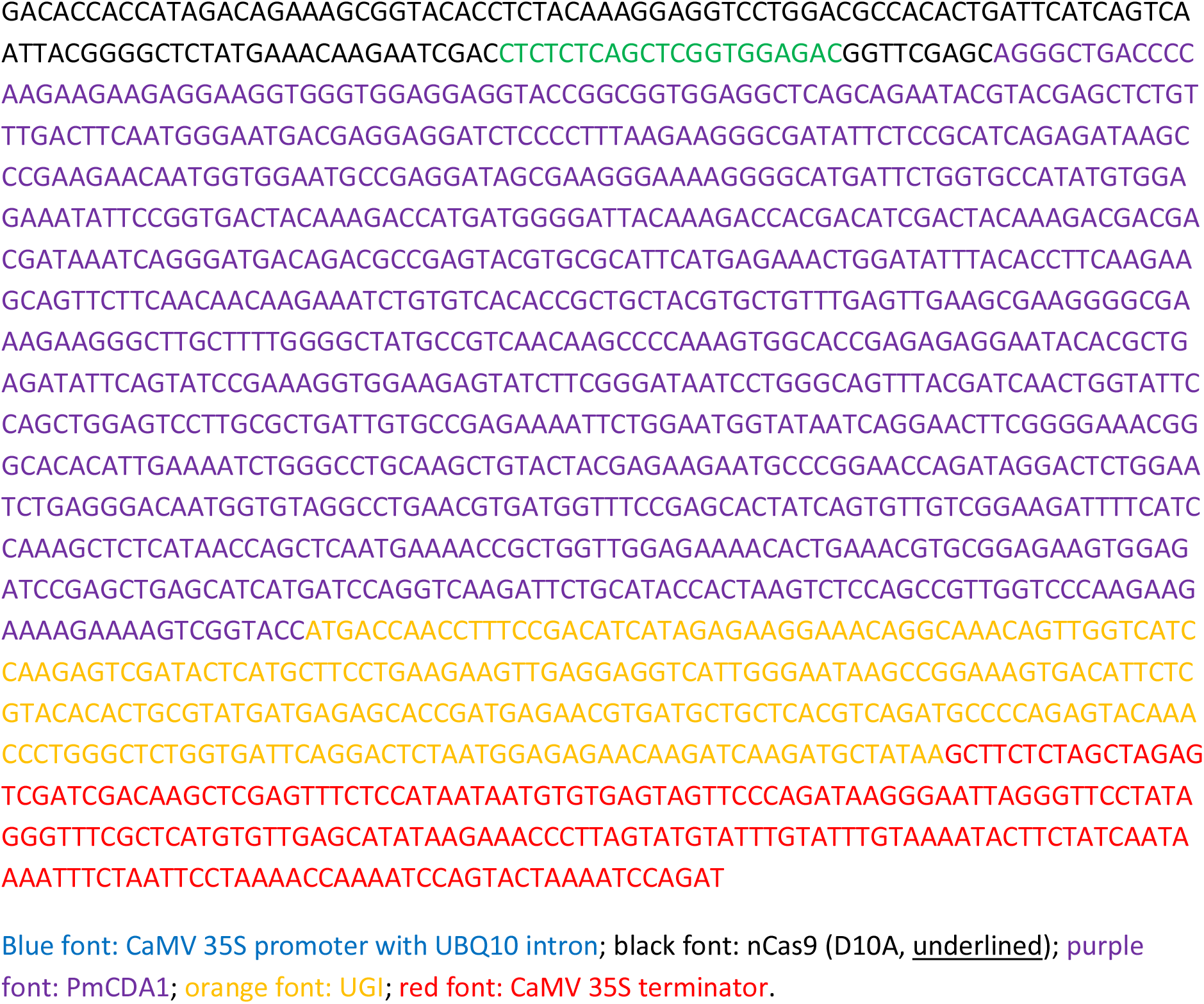
✥ **p35SI-nCas9-RT-t35S**

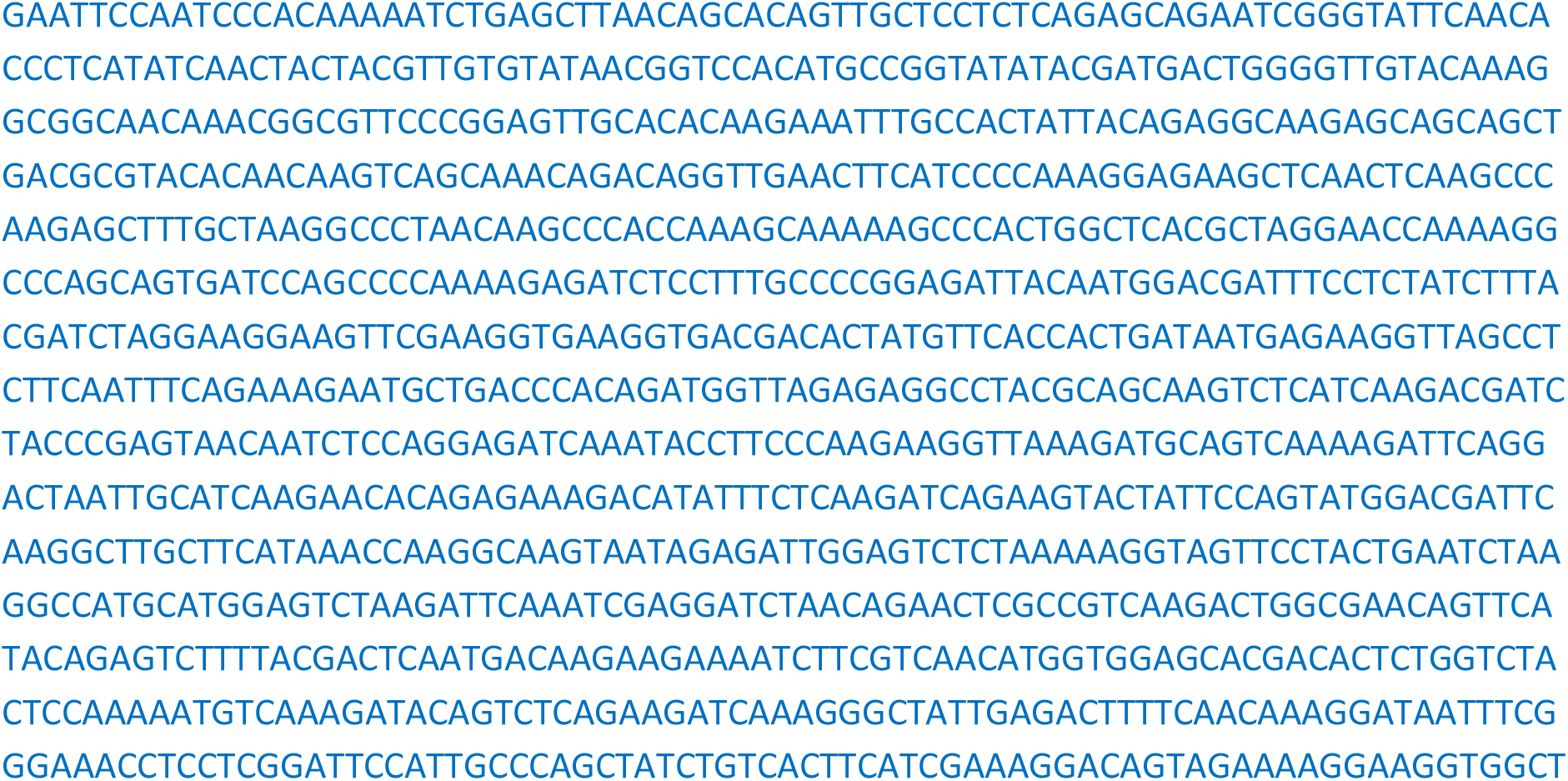

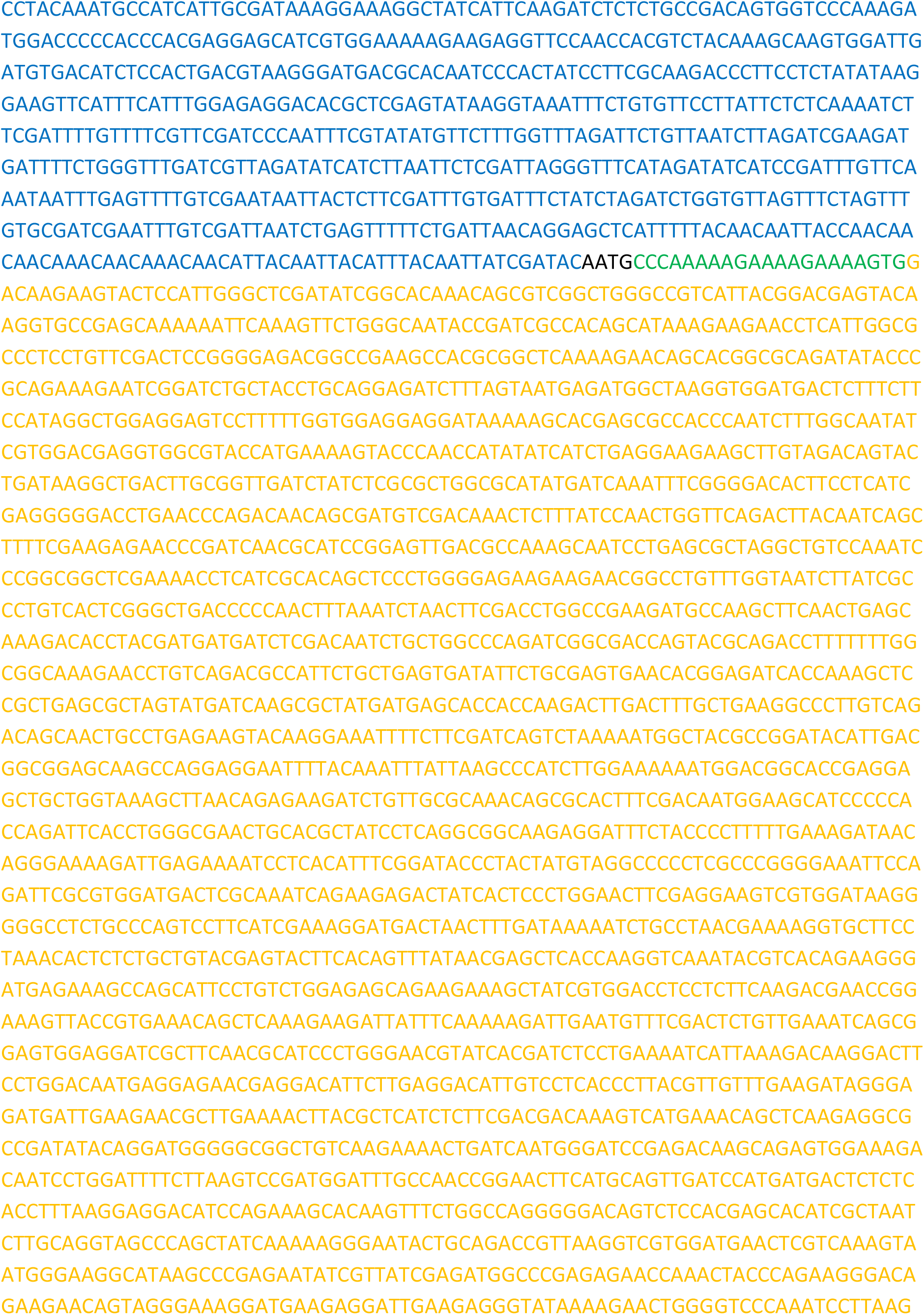

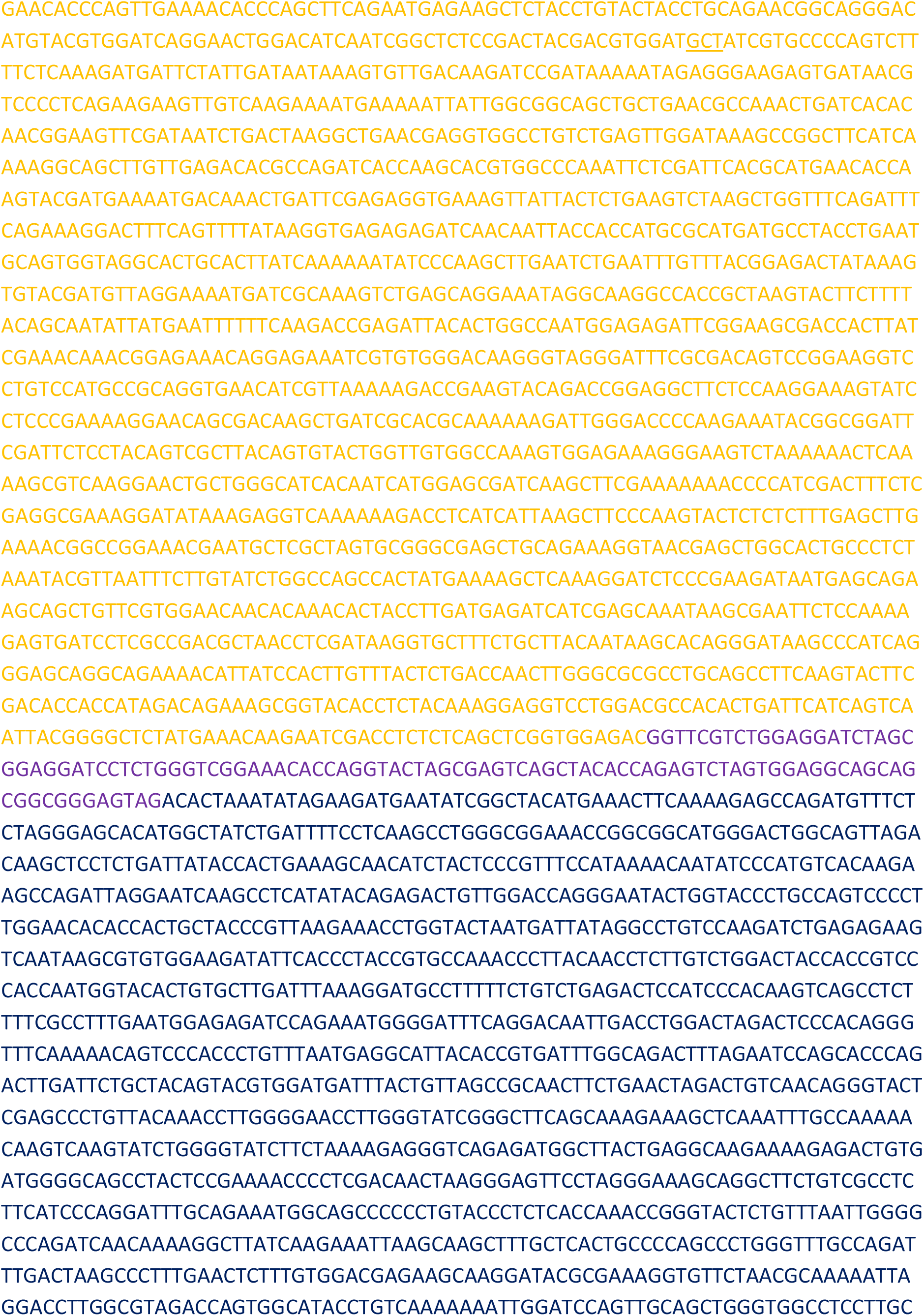

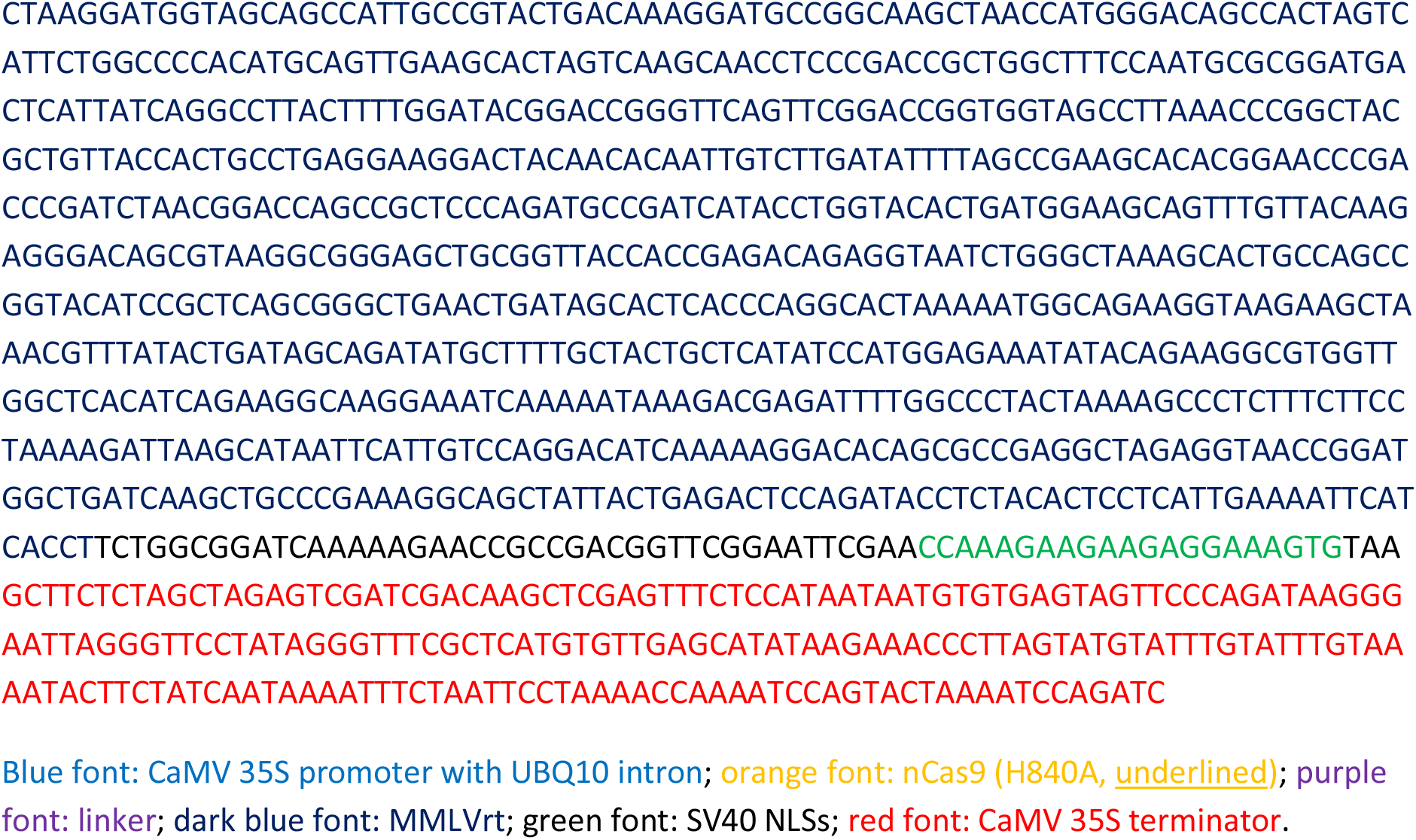
✥ **pegRNA1_CA**

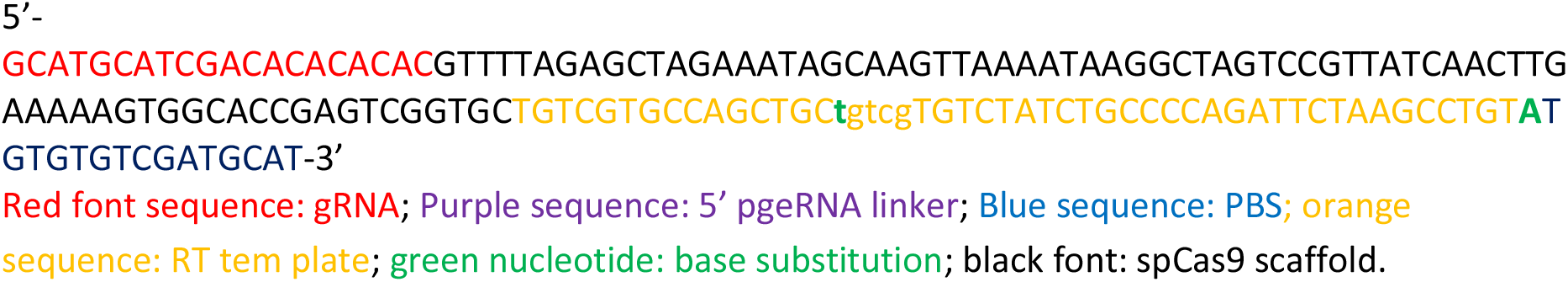
✥ **sgRNA_nick for second nick on the (CA)n substrate**

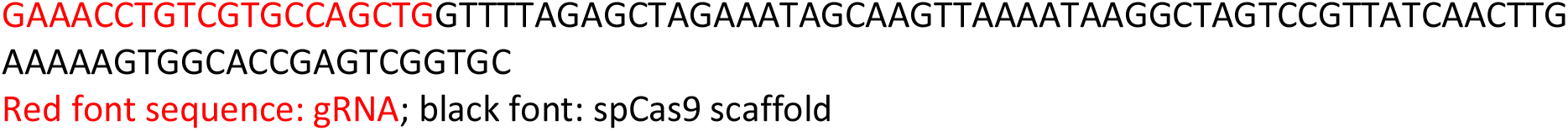
✥ **sgRNA_CA for base editing on the (CA)n substrate**

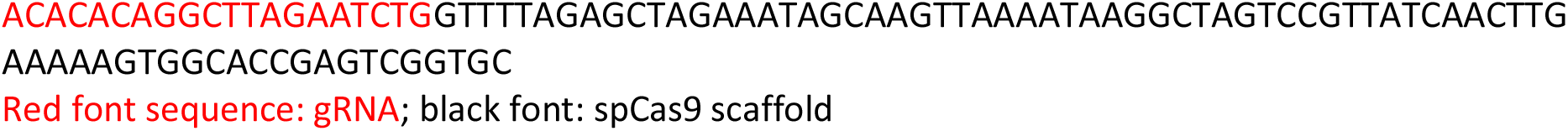
✥ **pegR_HKT12**

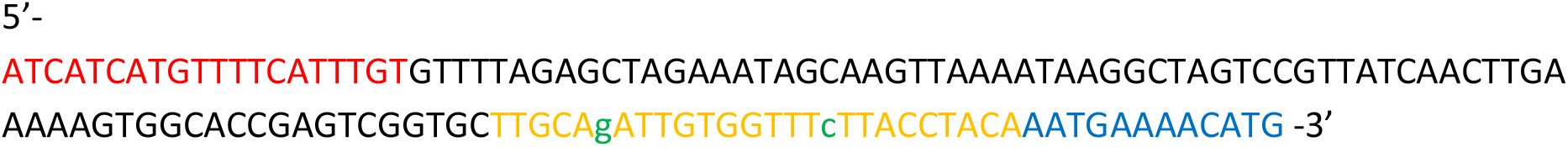

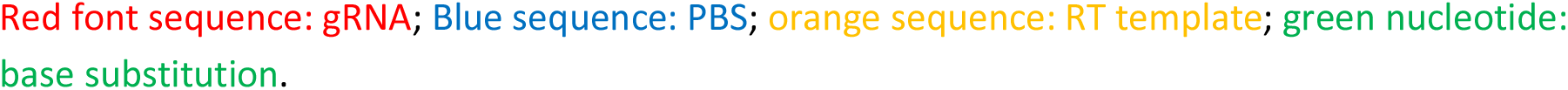
✥ **sgR_HKT12_2n for second nick at the SlHKT1;2**

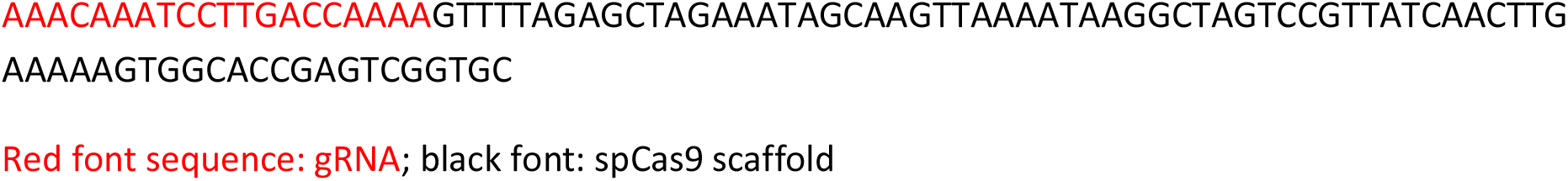
✥ **pegR_EPSPS1**

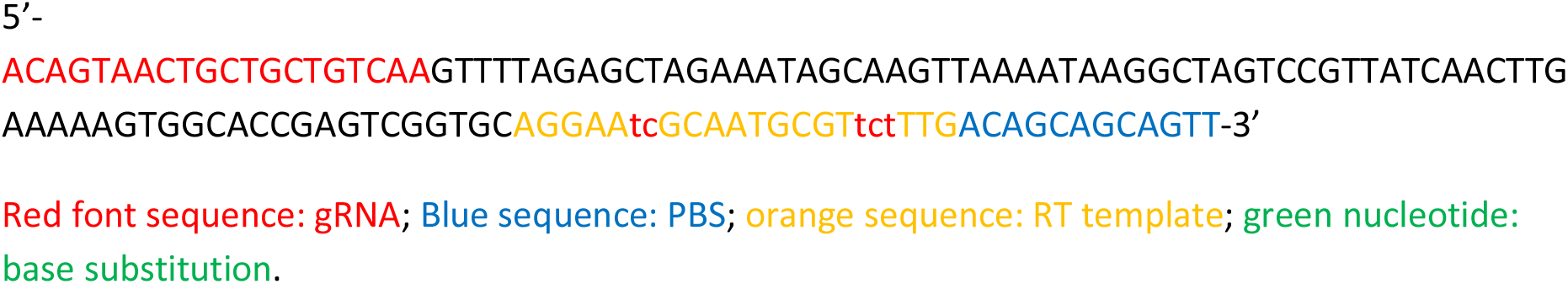
✥ **sgR_EPSPS1_2n for second nick at the SlEPSPS1**

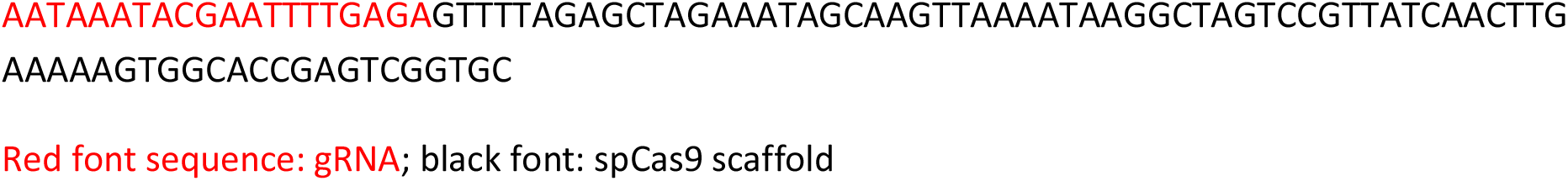
✥ **pegR_Or**

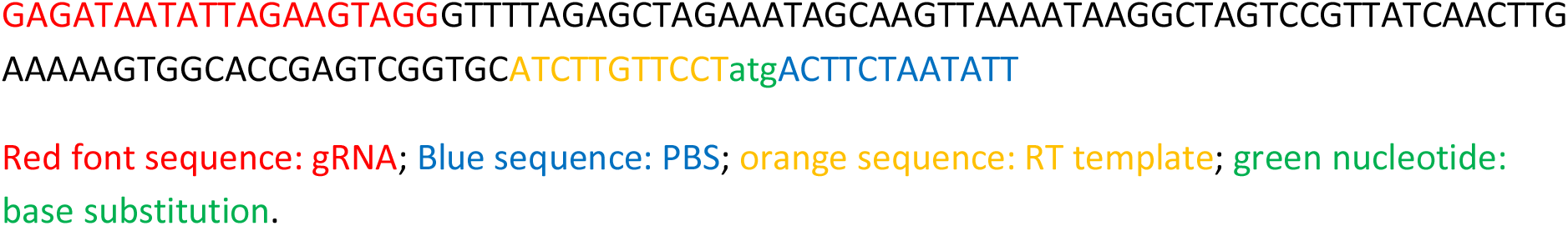
✥ **sgR_Or_2n for second nick at the SlOr**

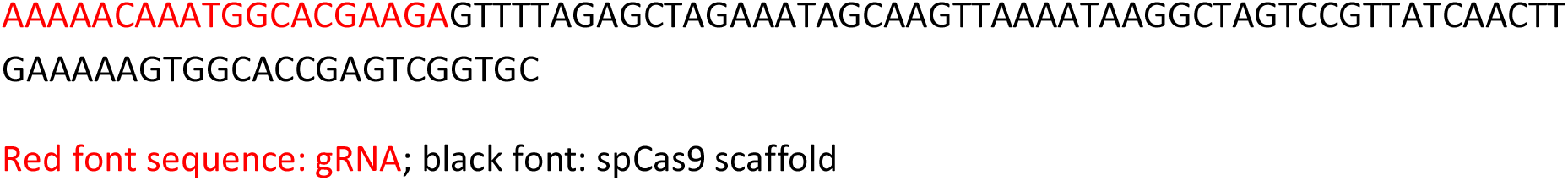
✥ **pegR1_ MBP21**

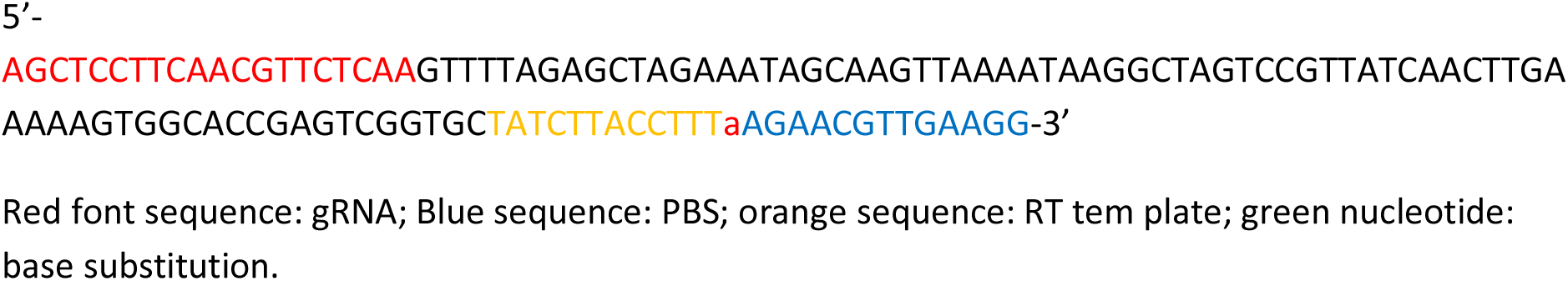
✥ **pegR1_WH9**

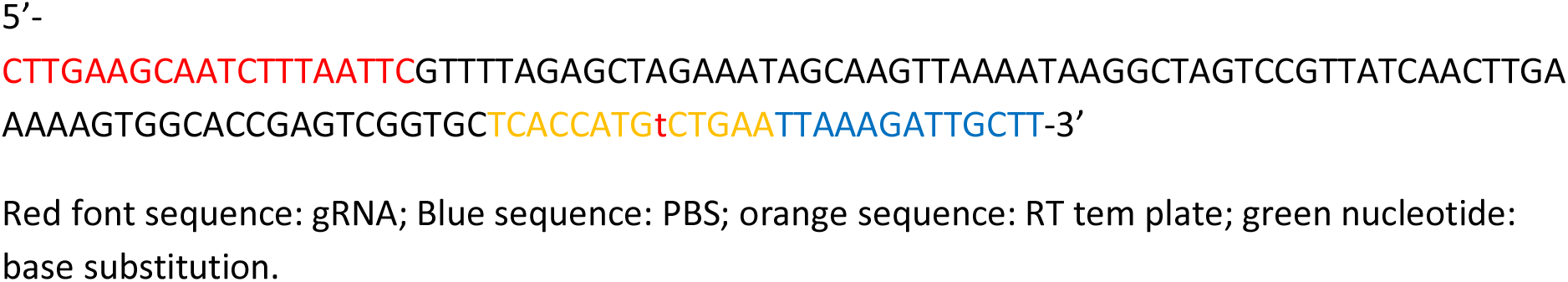
✥ **pegR1_KD1**

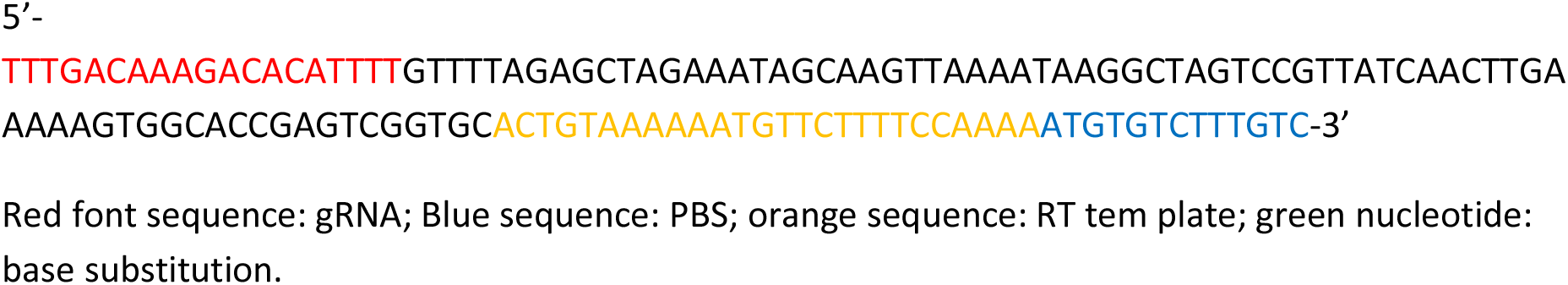
✥ **pegR1_PRD**

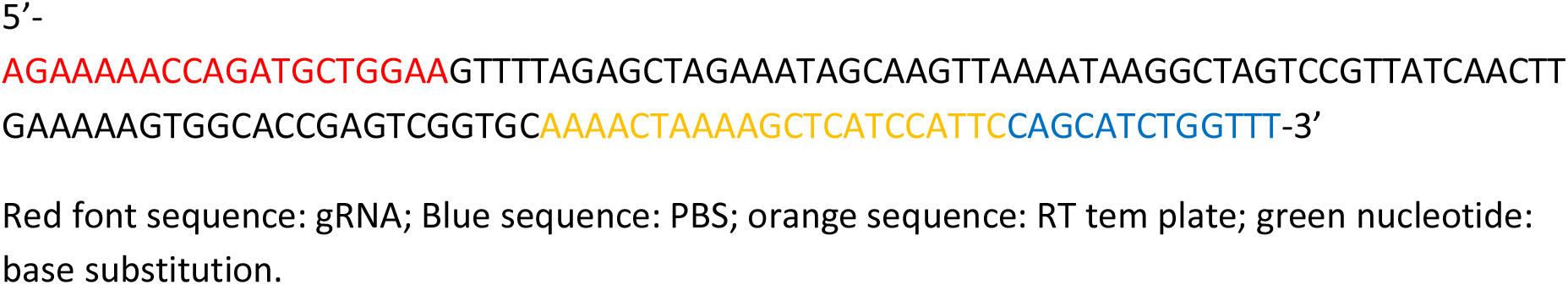
✥ **pegR1_ALC**

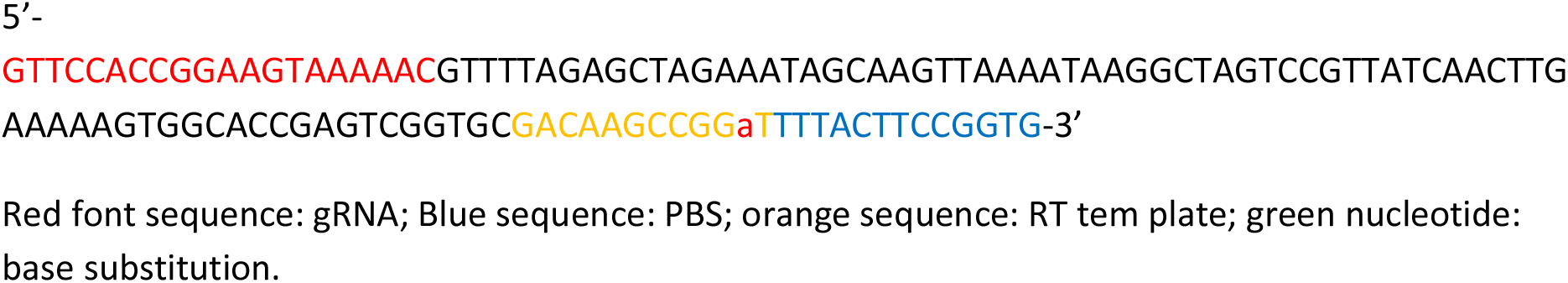
✥ **pegR1_DMR6**

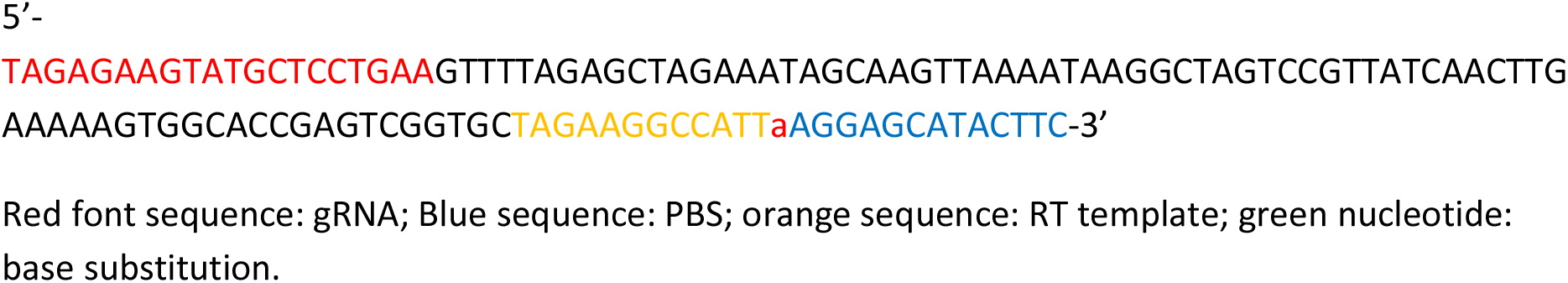
✥ **pegR1_ALS1**

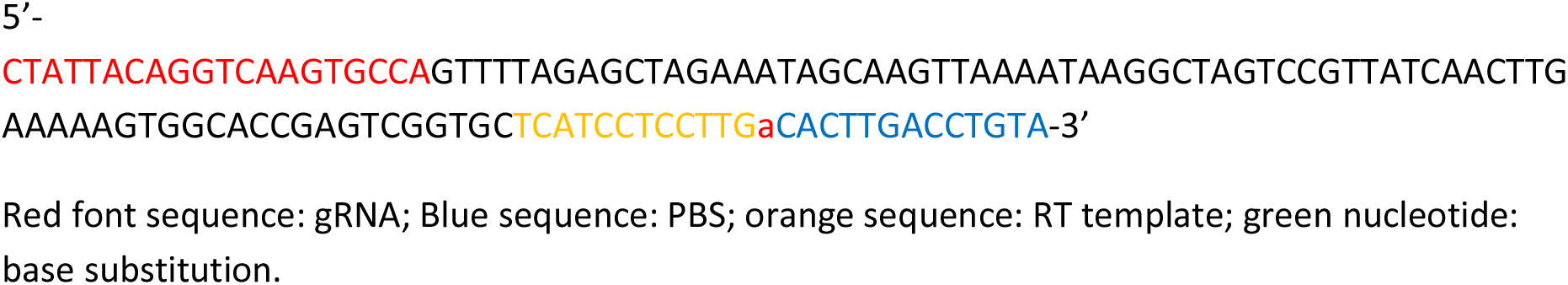
✥ **sgR1_ MBP21**

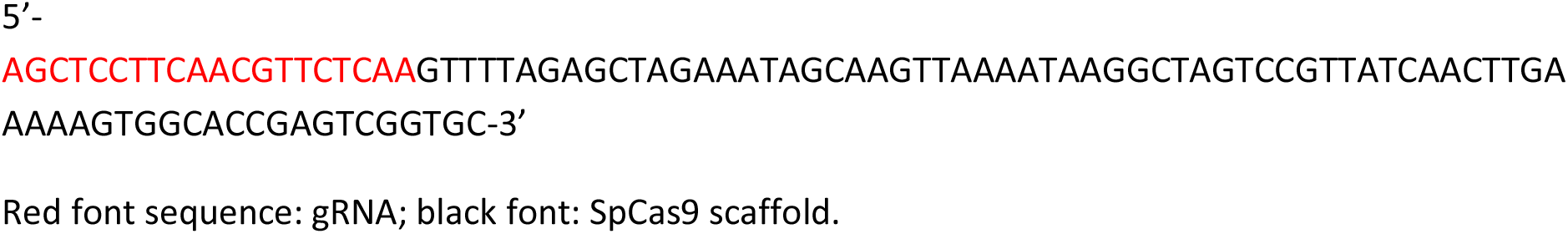
✥ **sgR1_WH9**

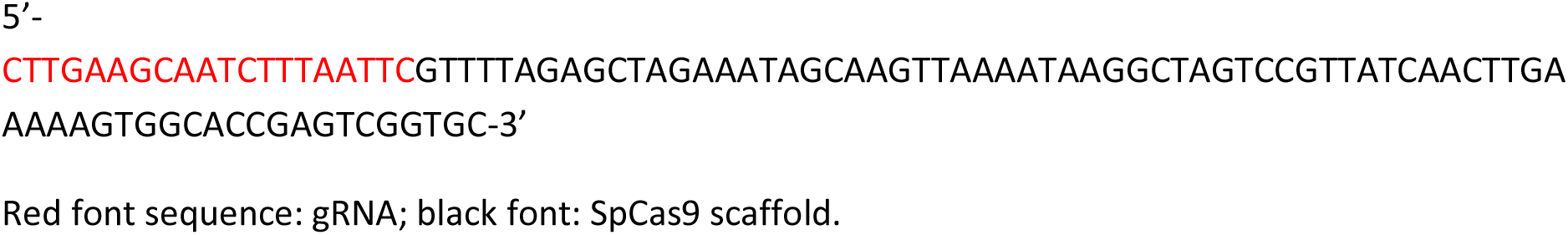
✥ **sgR1_KD1**

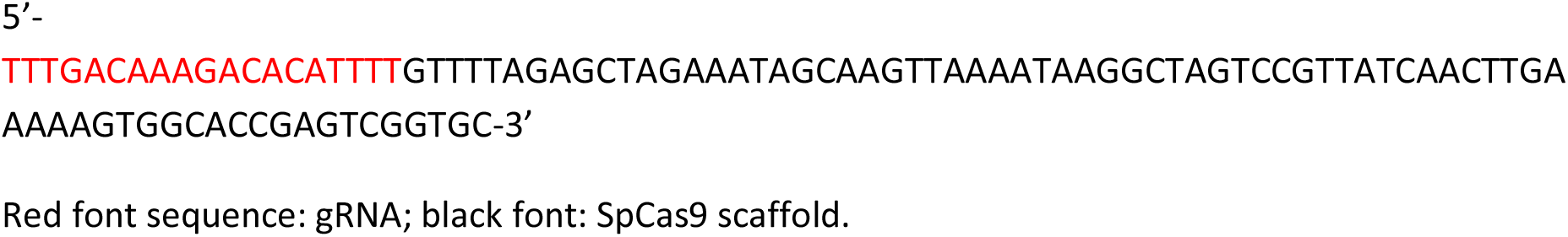
✥ **sgR1_PRD**

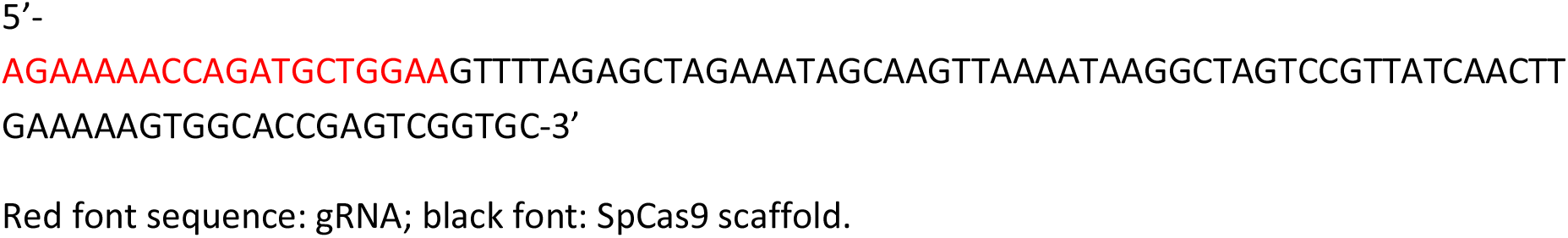
✥ **sgR1_ALC**

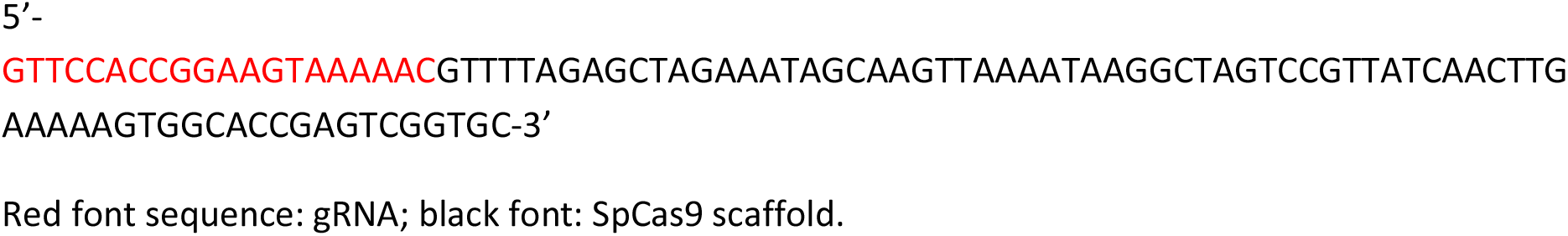
✥ **sgR1_DMR6**

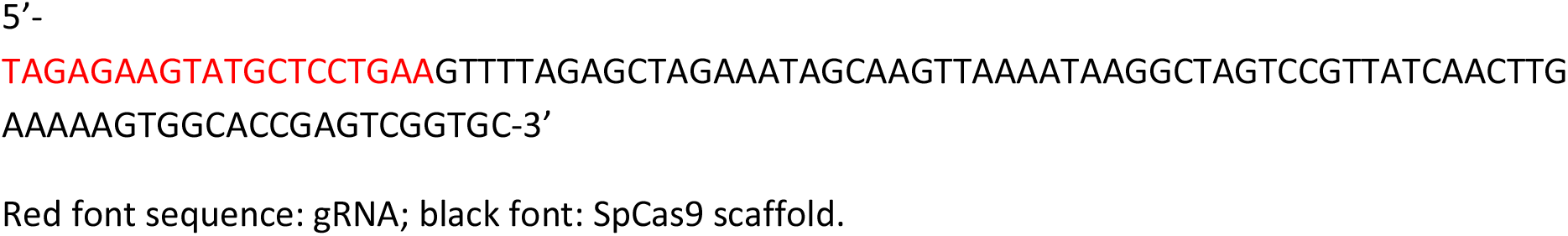
✥ **sgR1_ALS1**

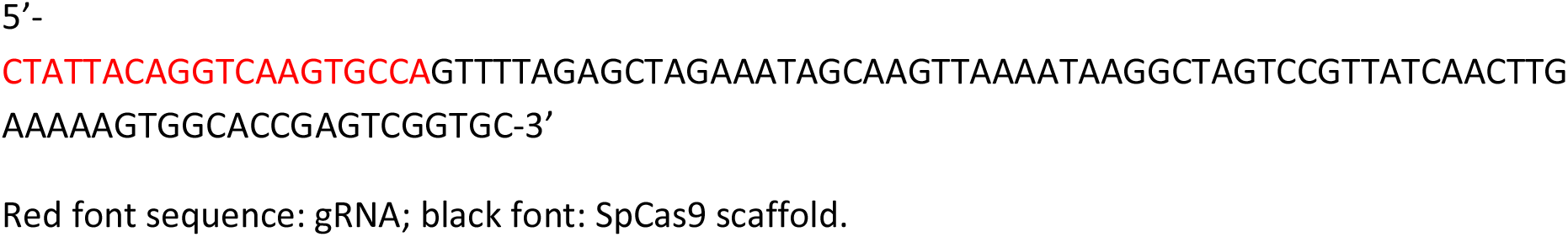
✥ **p35SI-Cas9-RT-t35S**

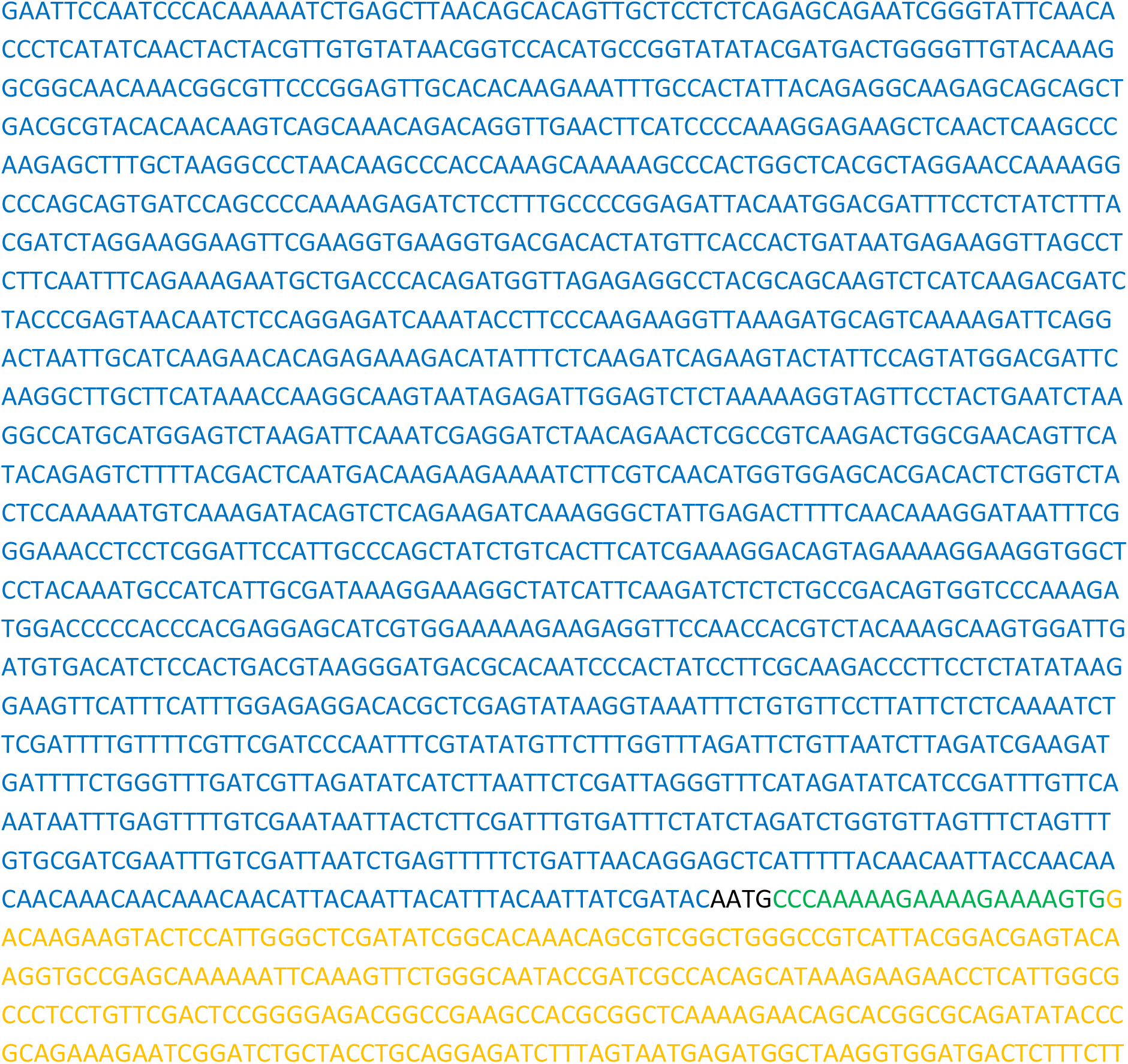

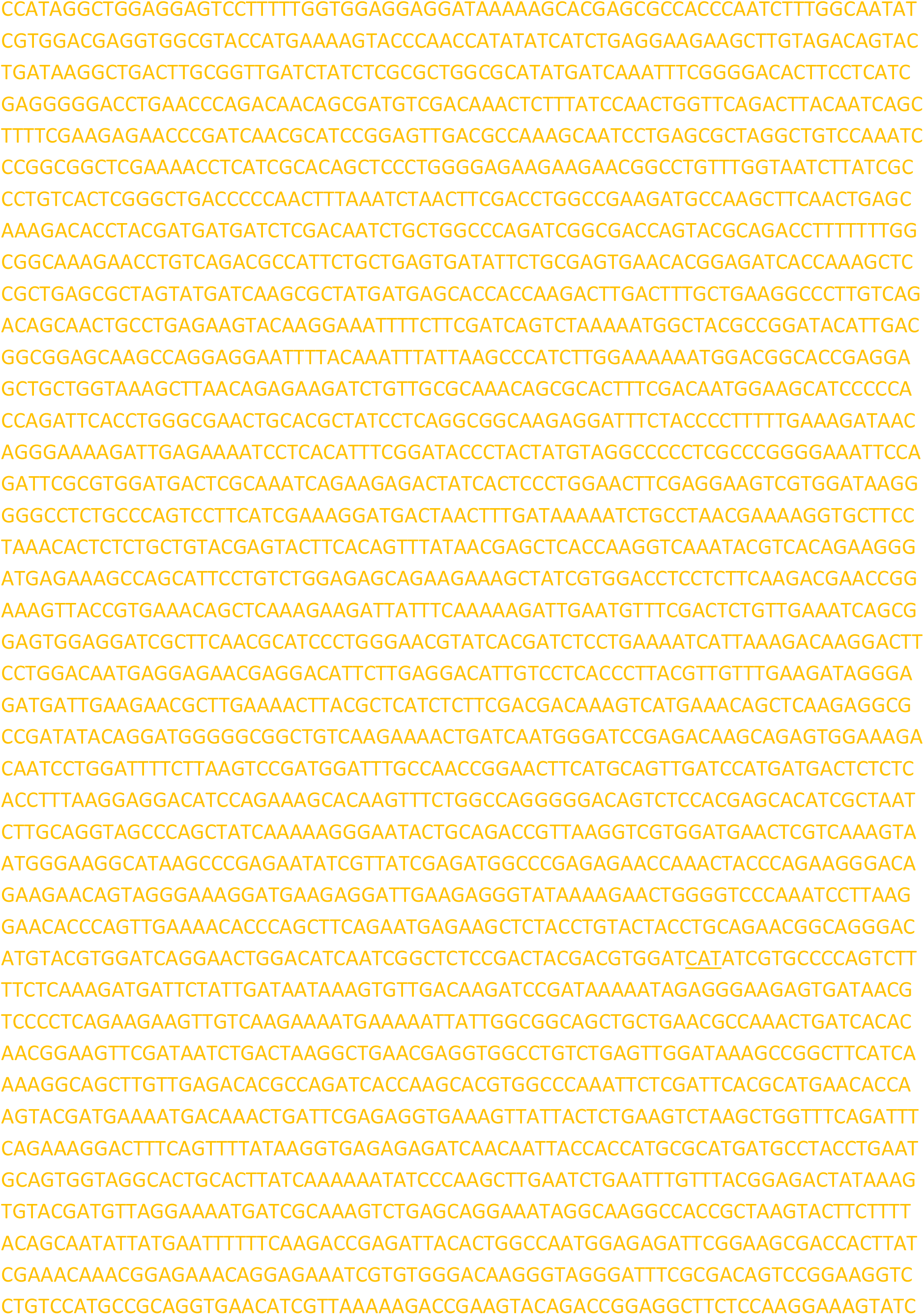

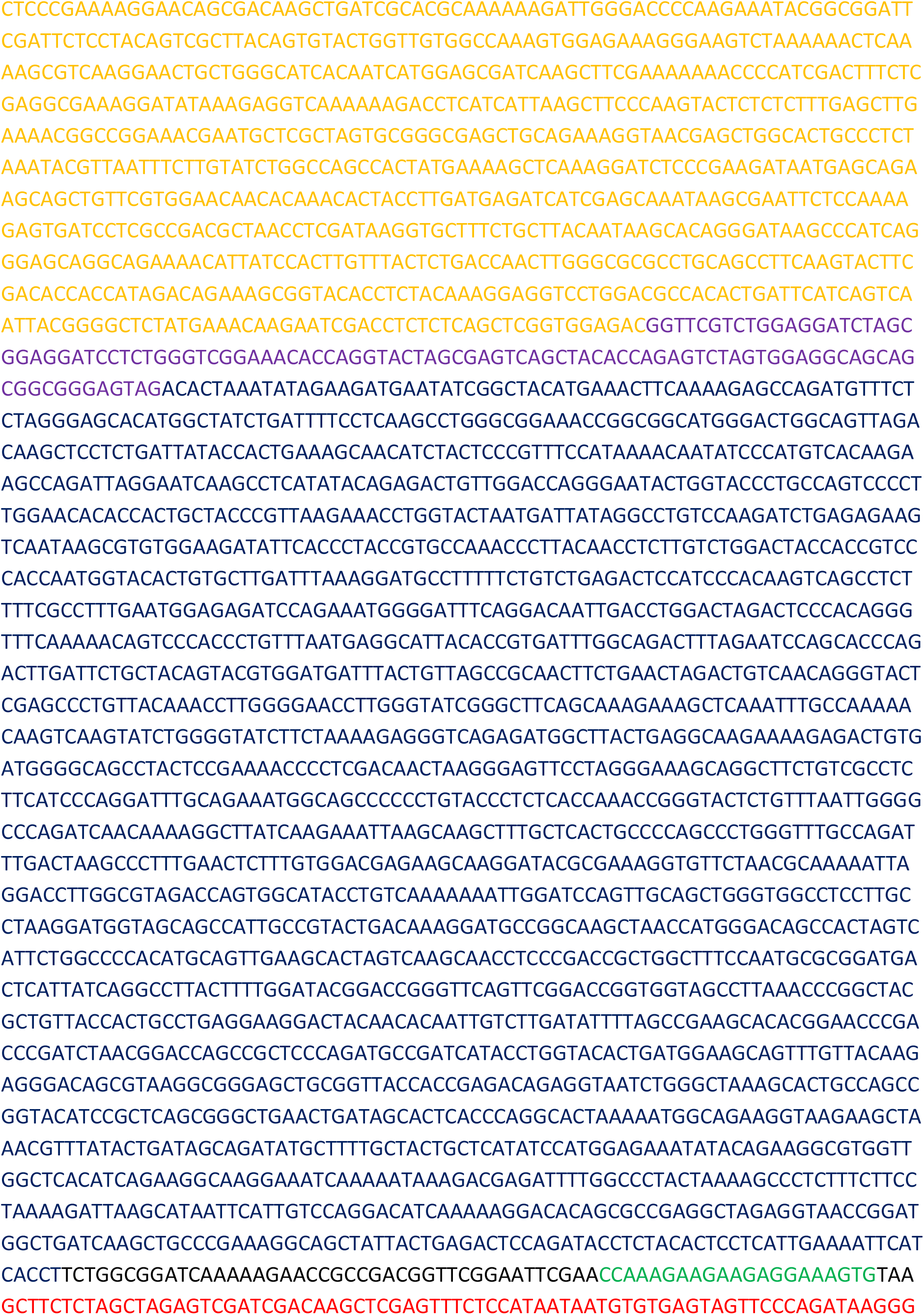

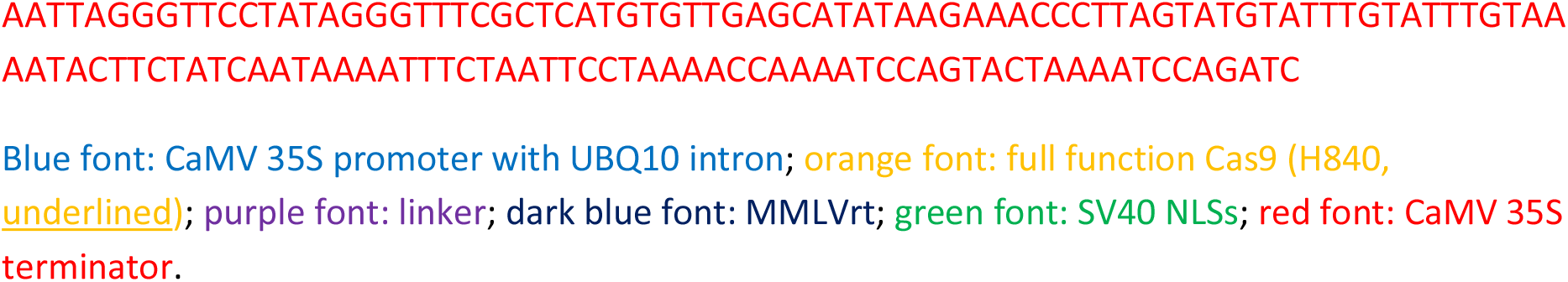
✥ **p35SI-Cas9-t35S**

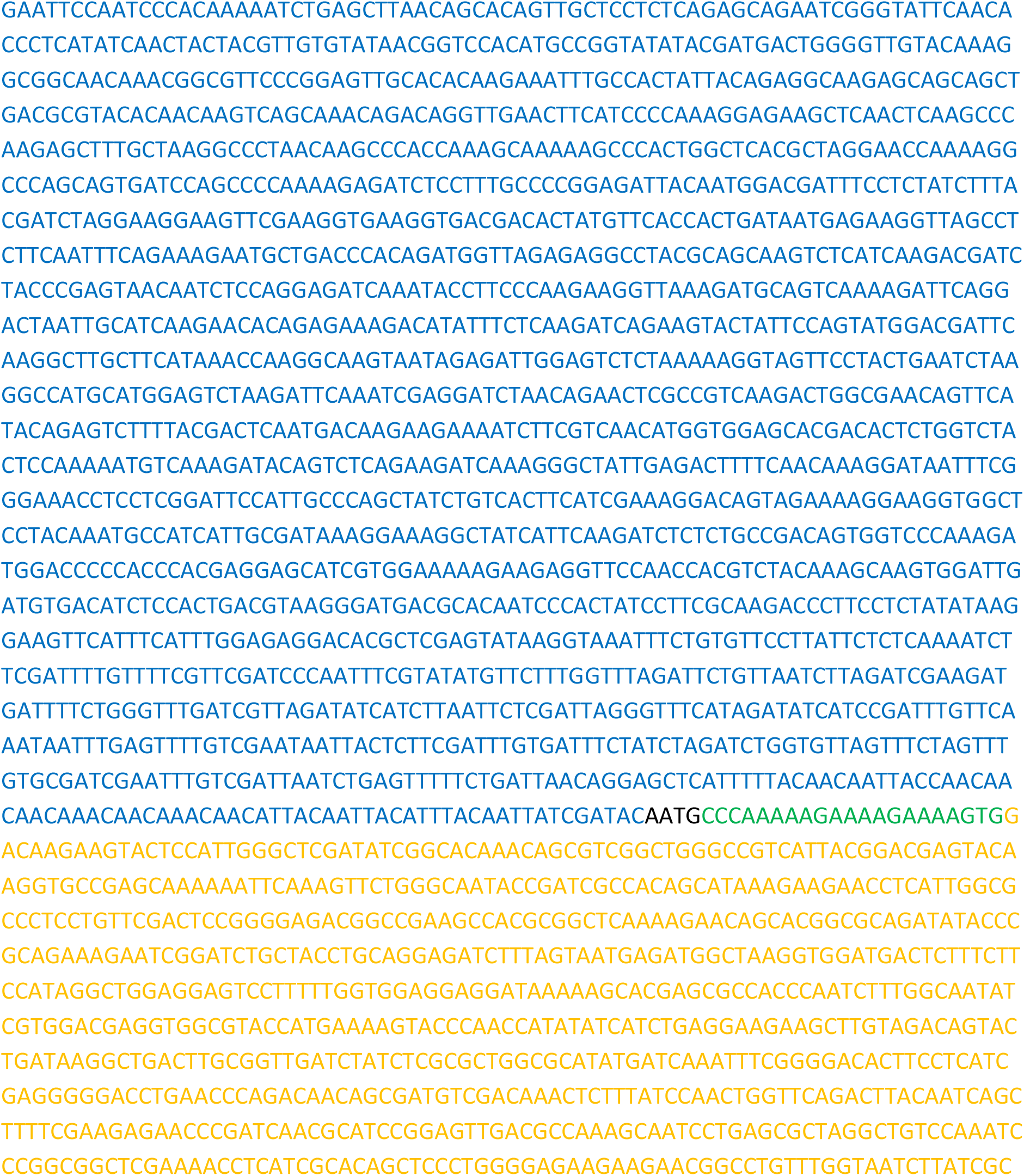

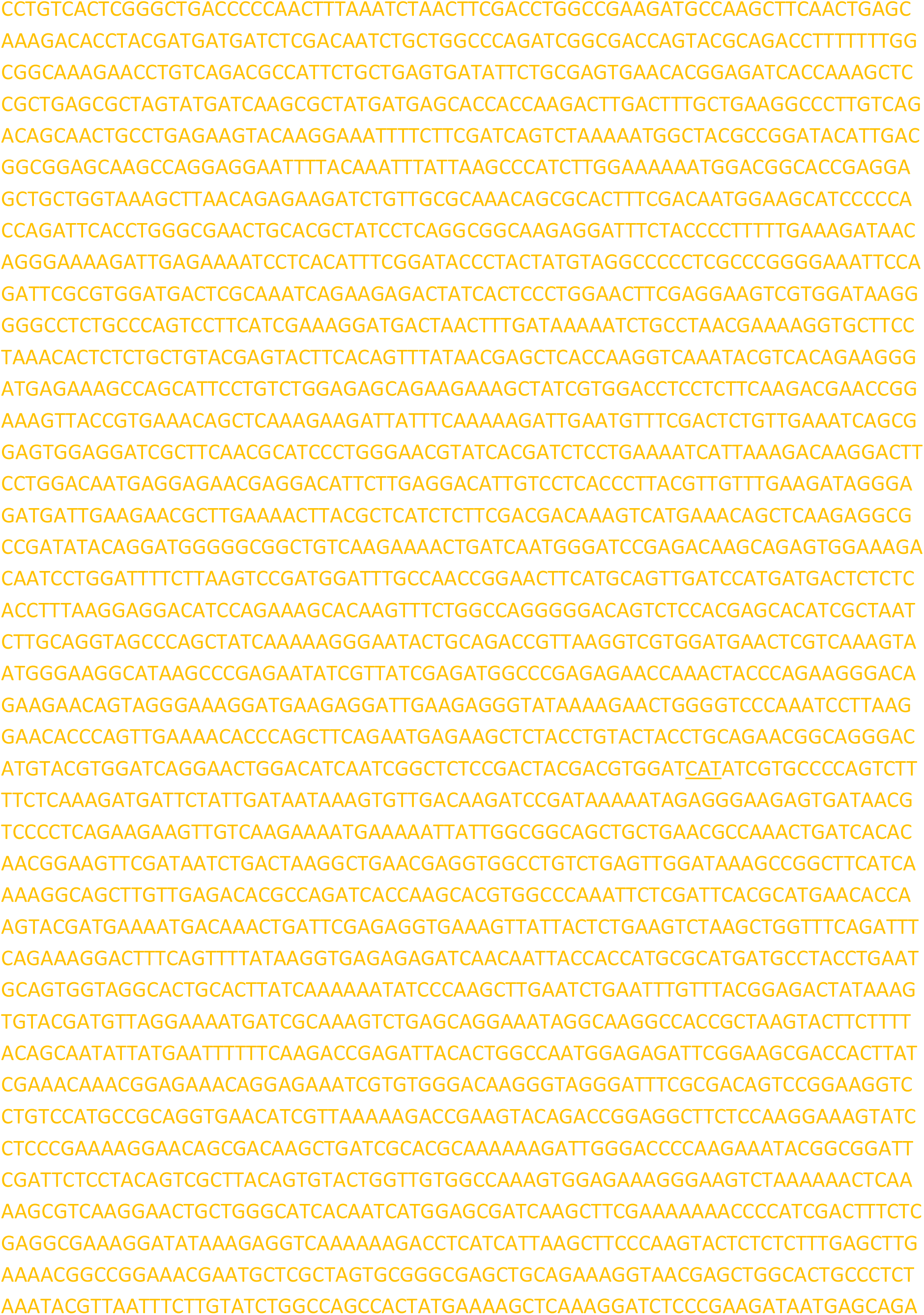

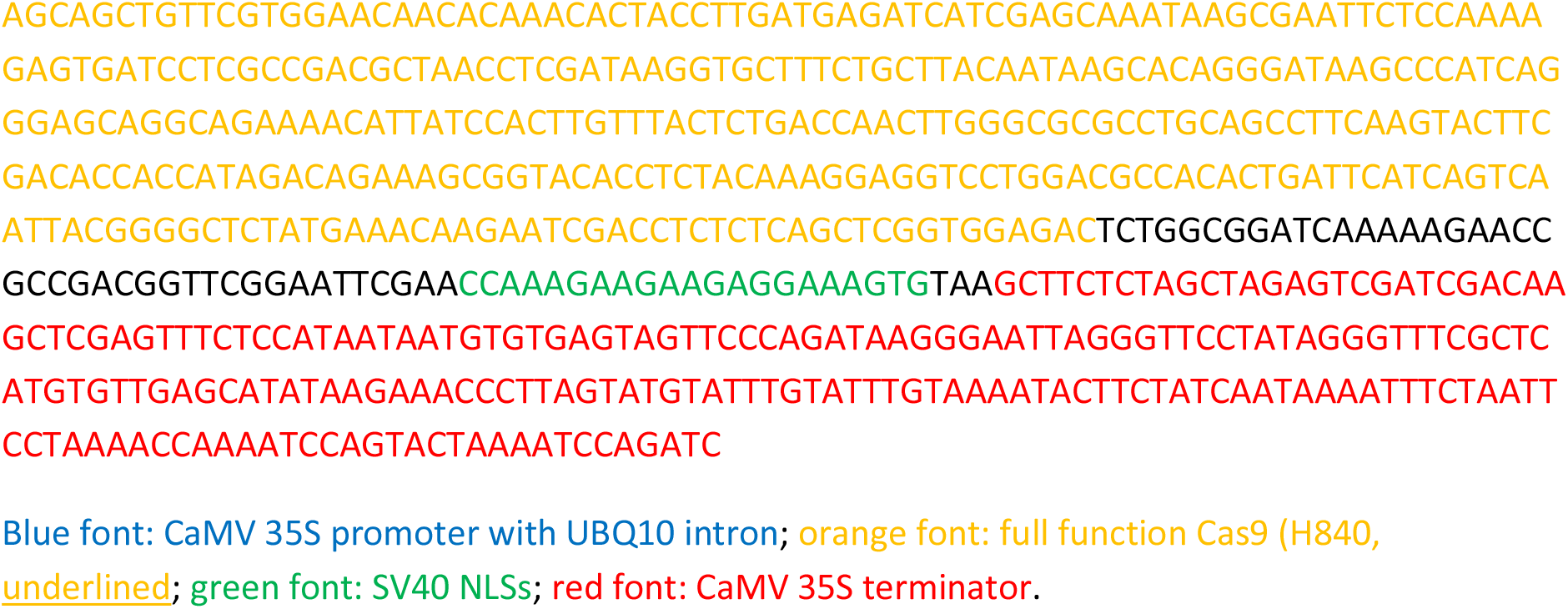
✥ **p35SI-nCas9-PPE-t35S expression cassette**:

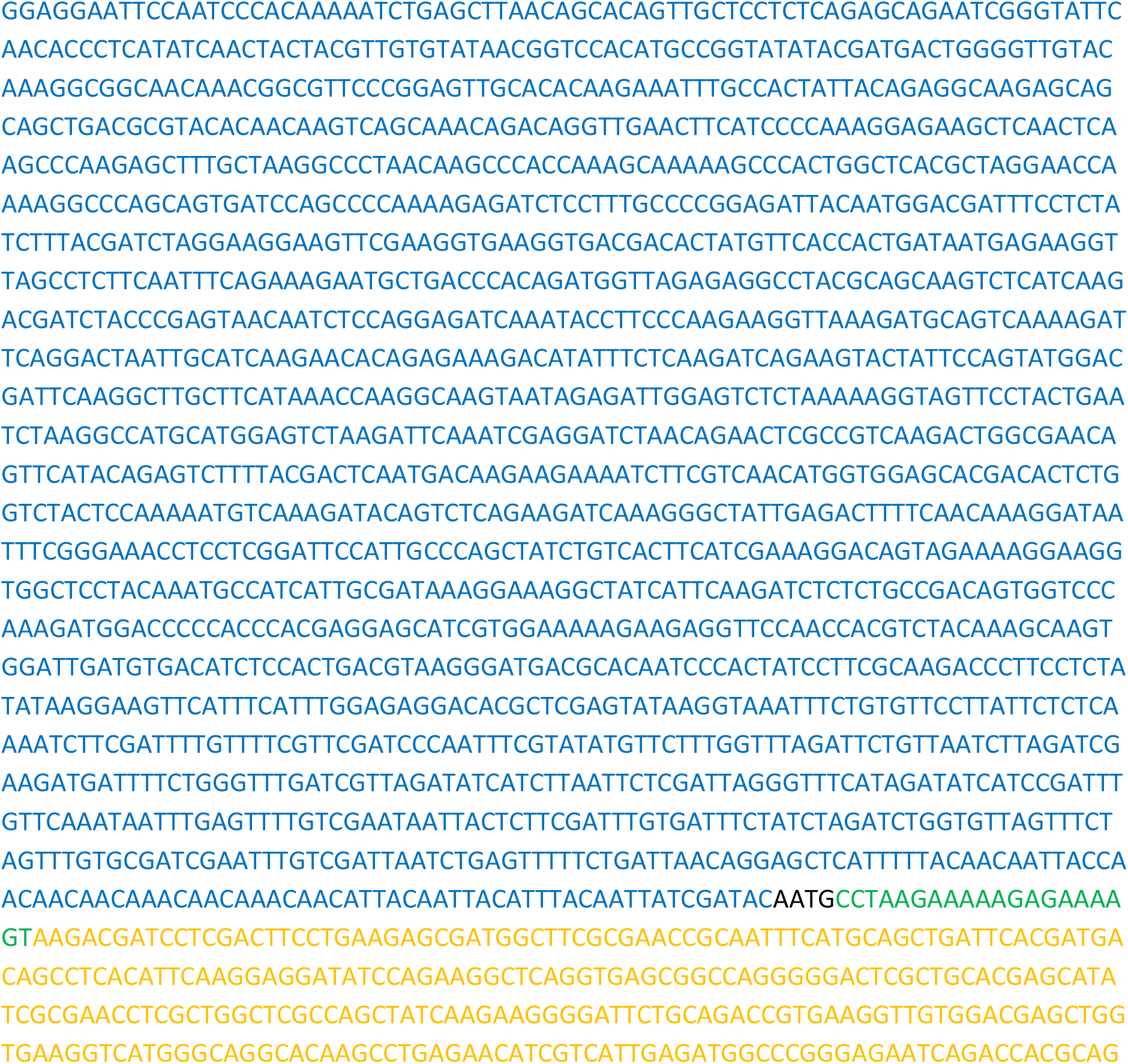

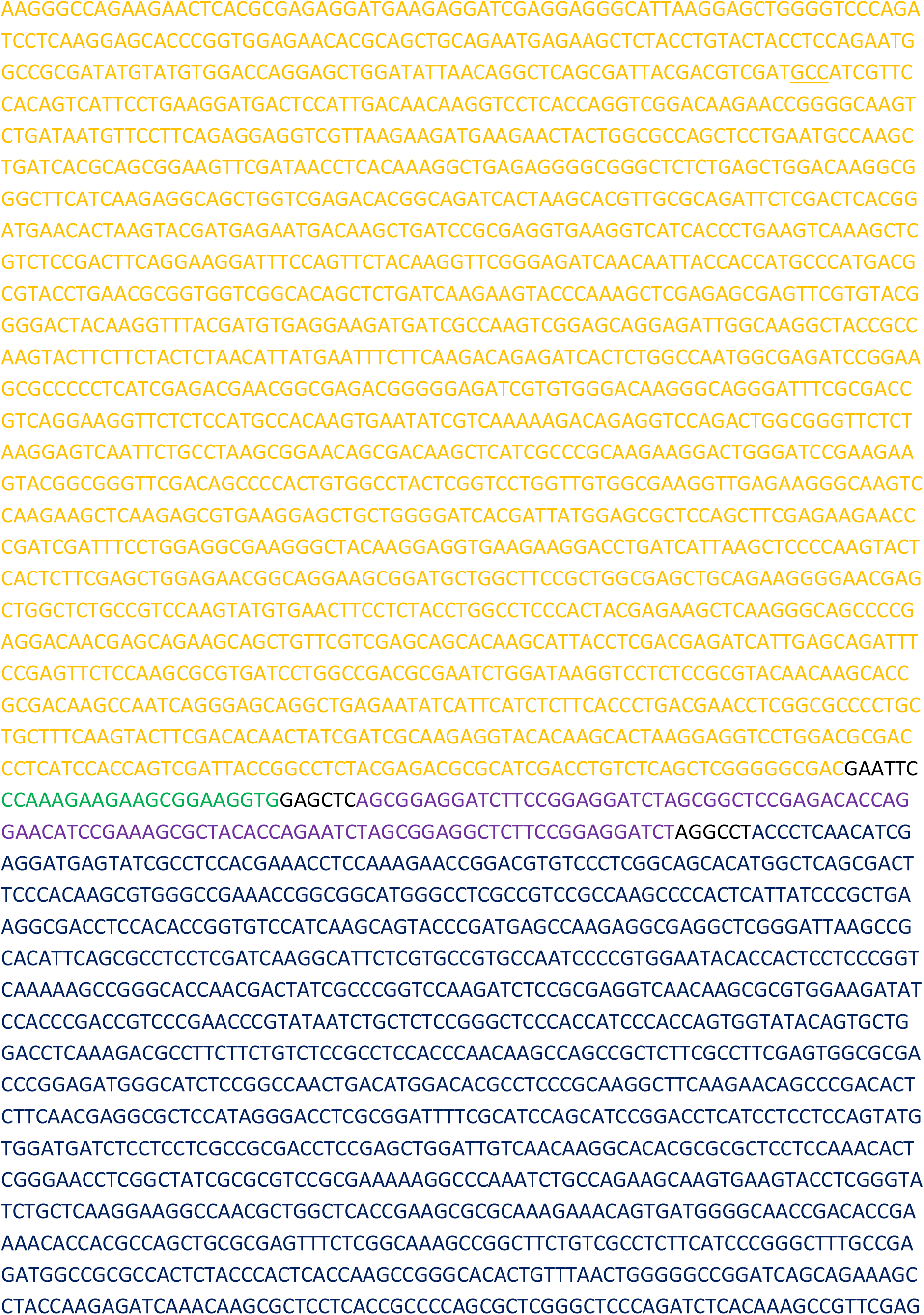

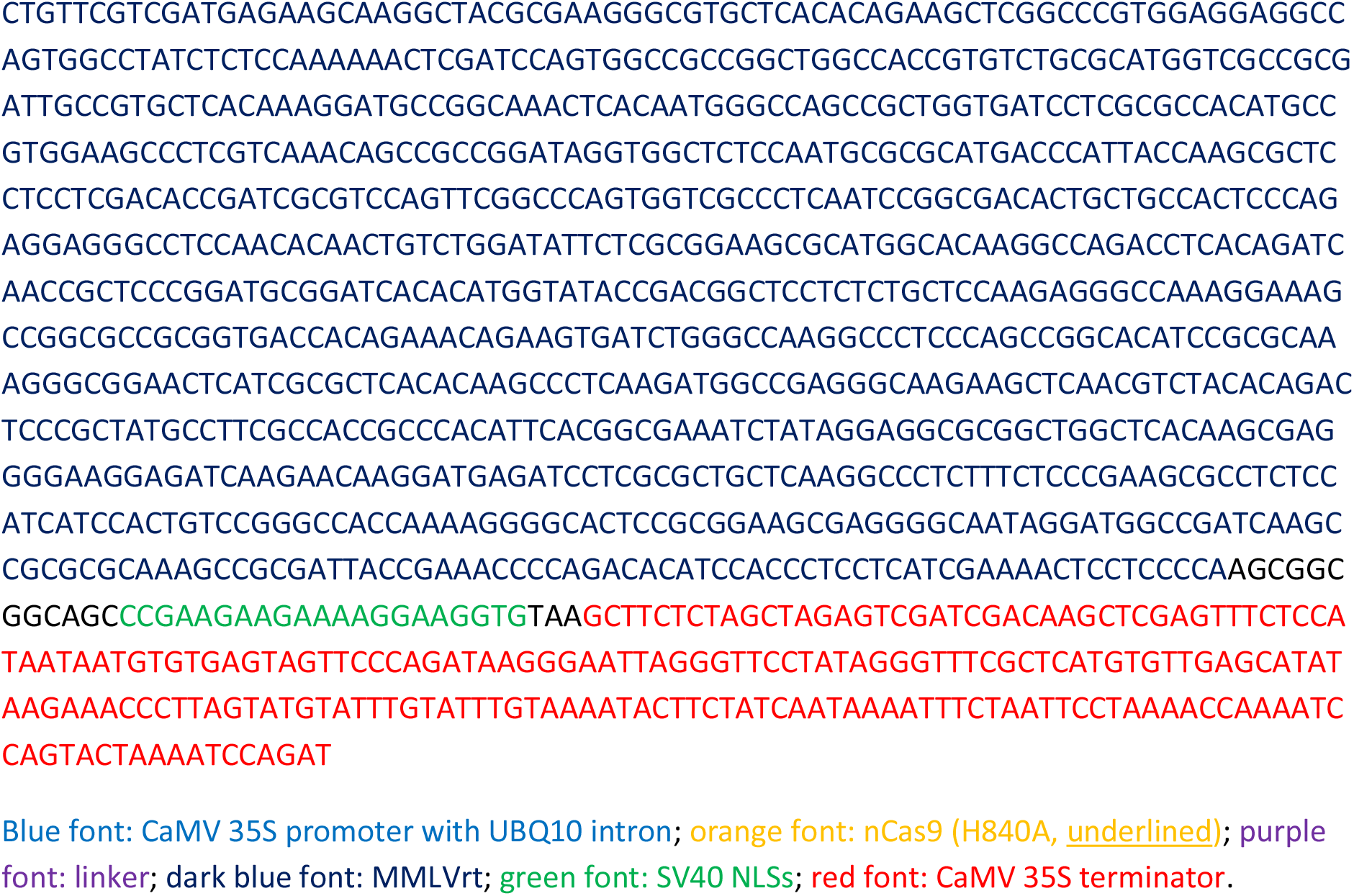
✥ **Plant selection marker**: pNOS-NptII-tOCS cloned from pICSL11024 (pICH47732::NOSp-NPTII-OCST) (Addgene Plasmid #51144).

